# Advanced Open-Source Experimental-Design Tools for Microplate-Based Assays with Acoustic Liquid Handling

**DOI:** 10.64898/2026.07.05.735934

**Authors:** Varunya M. Kattunga, Steven A. Wrobel, Chad A. Lerner, Victor M. Derycz, Elizabeth B. Stephens, Ian S. Brown, Hao Cheng, Sima Taghizadeh, Josef Byrne, Susan Gross, Susan Schneider, Chatura Senadheera, Asia Davis-Castillo, Shane Vistalli-Alvarado, Elena Goncharova, John C. Newman, Brianna J. Stubbs, Simon Melov, Gordon Lithgow, Lisa M. Ellerby, Julie K. Andersen, Akos A. Gerencser

## Abstract

Acoustic droplet ejection (ADE) enables nanoliter-scale liquid handling for complex microplate assays, yet translating experimental designs into validated, instrument-ready instructions remains a bottleneck. We present PickliPy, an open-source framework that converts spreadsheet-based assay designs into validated ADE picklists. PickliPy.Assay supports combinatorial, dose-response, and multi-addition time-course dispensing, while PickliPy.Screen extends to high-throughput workflows, including library reformatting and shortlisting. Across biological contexts, the framework generated reproducible, assay-ready plates and standardized execution in human cohort studies. Acoustic pre-dispensing deepened bioenergetic phenotyping of human skeletal muscle mitochondria, capturing substrate switching and sharpened dose-response precision in human pancreatic β-cells, revealing an age-associated change in succinate dehydrogenase kinetics. We benchmarked a wash-free, live-cell screen of mitochondrial function and morphology, in which deep-learning image analysis widened the assay window, and ADE enabled integrative dose-response co-response analysis. These tools, including their agentic use, make complex ADE experiments easier to design and scale from single benches to screening campaigns.

## Introduction

Acoustic droplet ejection (ADE) enables novel liquid dispensing strategies in scenarios ranging from focused academic research to discovery with high-throughput screening (HTS)^1,2^. ADE uses acoustic energy to transfer nanoliter-scale droplets in microplates from any source to any destination well without physical contact^3,4^. This allows library reformatting^5^, dose-response^6^, complex combinatorial designs^7^, compound pooling^8,9^, and lower vehicle concentrations than pipetting^10^. The current ADE technology applies to a variety of liquid types^11^, with well-demonstrated direct dispensing into cell cultures^10,12^ and making assay-ready plates for later use^13,14^. Here we focus on the translation of the complex experimental designs allowed by ADE from a human-readable format (spreadsheets) to a machine-executable series of instructions. This is a critical step in ADE and in HTS, where precise planning for thousands of liquid transfer events is routine. Contemporary, non-HTS lab bench research is also increasingly micro-scale and microplate-based. Below we describe openly available and generalizable software tools and their applications to improve both HTS and non-HTS experimental design.

The Beckman Coulter Echo ADE platform can execute simple mapped transfers and complex transfers based on protocols and picklists. The instrument can work stand-alone or as part of a robotic integration under the control of scheduling software. Picklists are tabular files specifying source wells, destination wells, and transfer volumes. The manufacturer’s commercial solutions such as Echo Plate Reformat, Echo Cherry Pick, Echo Combination Screen and Dose-Response are task-specific GUI applications for creating and executing transfer protocols. These apps optionally use picklists as input rather than creating them. Importantly, user-created picklists may be used in Plate::Works^TM^ (Revvity), Momentum (Thermo Fisher Scientific), and Green Button Go (Biosero). We provide design tools for picklists that can be used outside of the Beckman ecosystem.

Several alternative tools have been described to address aspects of ADE control, including the open-source FIMMCherry^15^ and PyEcho^16^ or software published as concepts, Echo QC^1^ (Genentech), Apothecary^7^ (AstraZeneca) and CherryPick^17^. Each of these alternative tools solved challenges for specific environments, including some tightly coupled with laboratory information management systems (LIMS) that may currently be a barrier in academic or start-up automation settings. At present, openly accessible ADE-support software remains fragmented: available tools either generate Echo-compatible transfer files for relatively constrained experimental designs, or implement specialized institutional workflows. We address the resulting practical gap by providing a locally deployable, publicly documented design-to-run-file workflow.

Here, we present PickliPy for the generation of Echo-compatible, validated picklists from spreadsheet-based assay designs. Developed for use in environments without LIMS, the picklist generation is based on a structured Excel workbook input, can be used with the common application DataWarrior^18^ or as an agentic tool. It produces outputs directly readable by Revvity Plate::Works^TM^ or other systems following the instrument’s API specification and generates metadata for downstream analysis. We demonstrate the utility of PickliPy across a range of experimental applications, including assay-ready plates for bioenergetics, flexible small-molecule library reformatting, dose-response, and combinatorial experiments.

## Results

### Two picklist paradigms: assay and screening

To translate intuitive plate maps into optimized and validated machine-executable series of instructions (picklists) and metadata files, we developed two complementary workflows to address major use cases: 1) PickliPy.Assay helps dispense single-plate tool libraries, creating combinations, dose-response series and time-course additions using multiple plate maps and optional randomization (Fig. 1a). 2) PickliPy.Screen helps library screening with reformatting and variable-dose dispensing where compounds in a multi-plate library are automatically assigned to wells of multiple assay (destination) plates using a single generic layout (Fig. 1b).

**Fig. 1.**
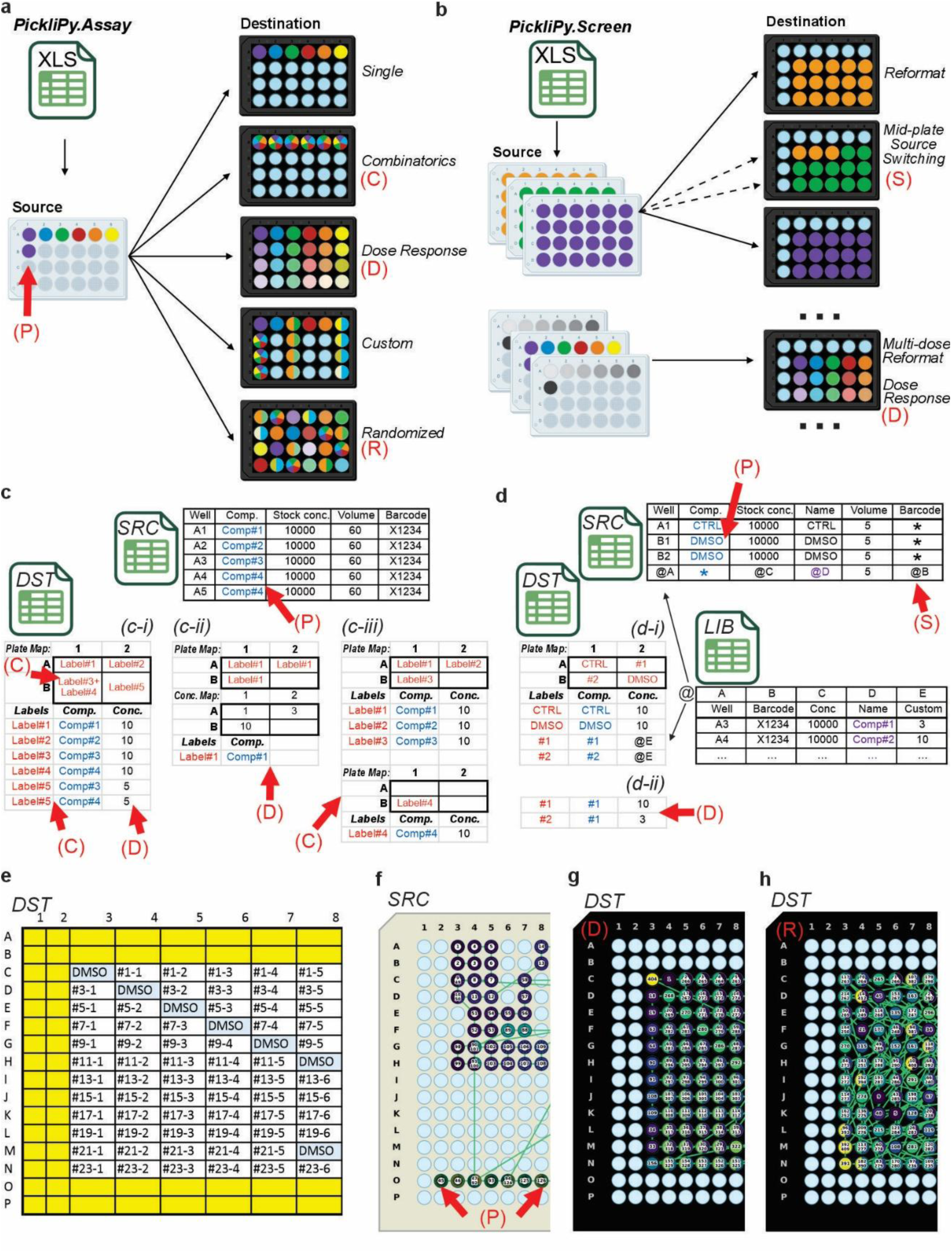
Spreadsheet designs generate executable ADE picklists for assay and screening workflows. **a)** PickliPy.Assay supports dispensing single-plate tool libraries creating combinations (arrow C), dose-responses (D) with optional randomization (R), and well pooling (P) using a single Microsoft Excel design document. **b)** PickliPy.Screen supports library screening with reformatting, adding controls, and using multiple doses per compound, advancing source and destination plates as needed (S). **c)** For PickliPy.Assay, the SRC worksheet defines source plate compound locations, names and stock concentrations. The DST worksheet defines one or more dispense plate maps, allowing users to specify compounds and their concentrations in multiple ways: **c-i)** one or more labels per worksheet cell [where labels (in red) are associated with compound stocks defined in SRC (in blue) and with final concentrations] or one or more compounds with concentration per label (bottom); **c-ii)** a single label per cell plus a concentration map to allow setting numeric values for concentrations and using autofill in Excel; and **c-iii)** multiple plate maps, including strategies shown in **c-i)** and **c-ii)** that are combined for dispense (see also Fig. S2). **d)** For PickliPy.Screen, the SRC worksheet defines controls added to source library plates and library compound locations referencing the LIB worksheet with a single wildcard row with a * as compound name. The DST worksheet defines only one generic plate map where any labels other than the controls defined in SRC are automatically populated from the LIB worksheet. This allows using **d-i)** explicit or referenced final concentrations from LIB worksheet rows; and **ii)** multiple doses of the same library compound (see also Fig. S3). **e)** Dose-response dispense using the strategy illustrated in **d-ii**, showing a part of a 384-well plate design. Label prefixes #1 to #23 refer to 23 library compounds, while the label suffixes refer to different concentrations for each compound (see corresponding label definitions in Fig. S3). **f-h)** Visualization of plate movements based on the generated picklist for the SRC plate (**f**) corresponding to the design in (**e**). Numbers in wells indicate the order of dispensing. (P) indicates compound well pooling for DMSO. **g)** Well-to-well plate movements are optimized for the shortest path in the DST plate corresponding to **e)**. **h)** Well-to-well plate movements using the design in **e**) with randomization PickliPy.Assay.

Both methods use a single design file as input, containing an inventory or source (SRC) worksheet (Fig. 1c, d) defining source plate compounds and a destination (DST) worksheet defining dispense positions and concentrations. The two methods differ in how these worksheets are handled, providing explicit dispense patterns for PickliPy.Assay (Fig. 1c-i, c-ii, c-iii) or high-throughput generalization with an auto-filled layout (Fig. 1d-i, d-ii, e) from a third, library (LIB; Fig. 1d) worksheet for PickliPy.Screen. Expected source well volumes are tracked during dispense planning, enabling source well pooling to avoid depletion (Fig. 1c, d, f; arrow p). Dispense order is optimized to minimize plate movements and is optionally randomized (Fig. 1f-h). The companion tools, PickliPy.QC and PickliPy.Visualize, enable post-execution quality control by parsing instrument log files and illustrate plate movements, respectively (Fig. 1f-h).

Using a Beckman Coulter Echo 650 ADE device integrated into a robotic workcell (Fig. S1), we first demonstrated acoustic dispensing of a pattern for dot-blots using a single plate map and label definitions for dispense volumes (Fig. 2a-b).

**Fig. 2.**
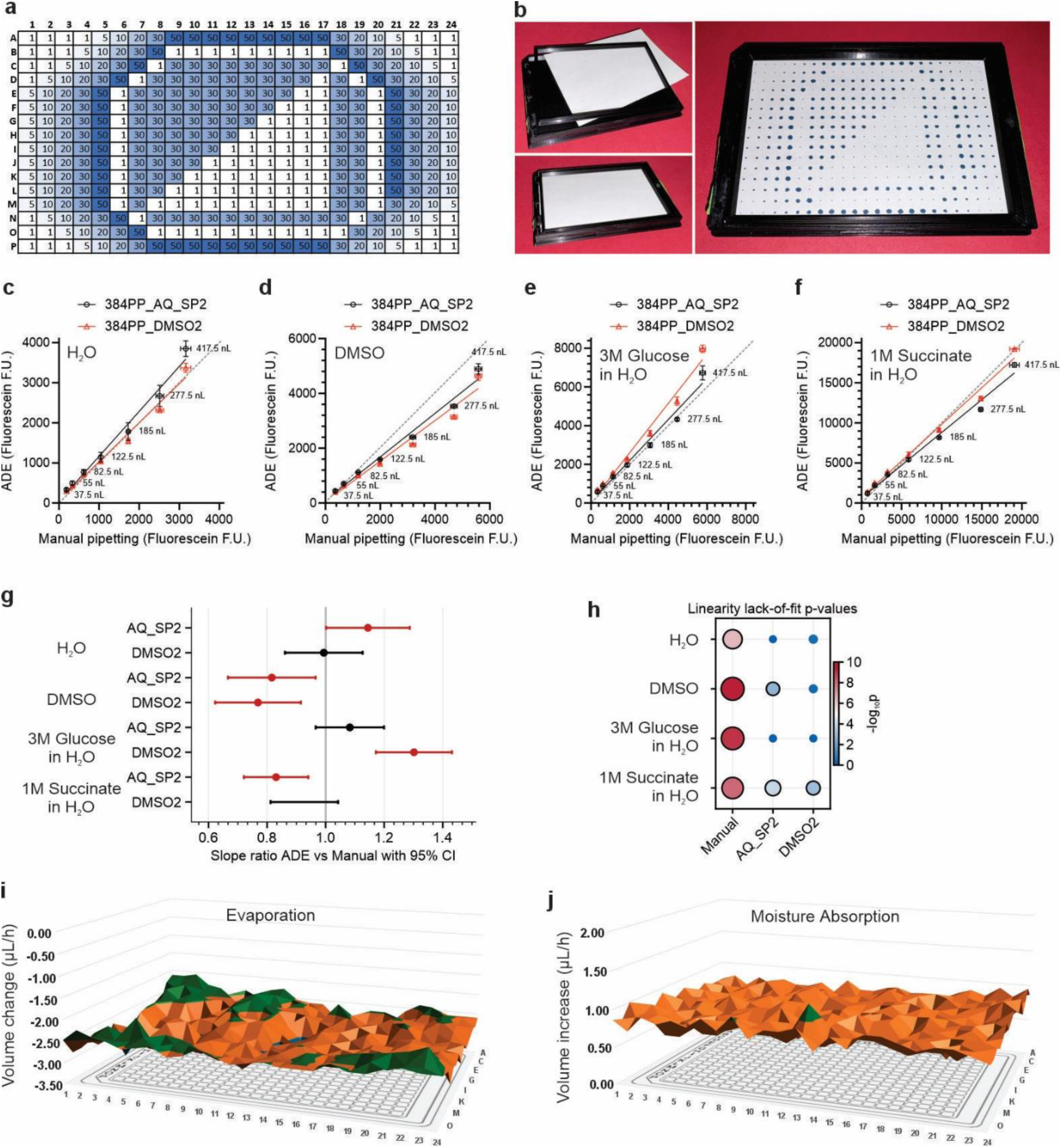
Protocol tolerance and source-plate residence define operating limits for ADE. **a)** Concentration map for dispensing a pattern with PickliPy.Assay. **b)** Custom 3D-printed microplate adaptor for blotting paper with ADE-dispensed pattern corresponding to **a**). **c-f)** Comparison of well-to-well variations of ADE to manual pipetting using fluorescein as tracer, dispensing H_2_O (**c**), DMSO (**d**), and aqueous solutions of glucose (3 M; **e**) and succinate (1 M; **f**). Each liquid was dispensed by an ADE protocol for aqueous liquids (384PP_AQ_SP2; black) and for DMSO (384PP_DMSO2; red). Data are mean ± SD of n = 3 well replicates, averaged from two experimental repeats. **g**) Comparison of manual and ADE dispense corresponding to **(c-f)** using unweighted linear-regression slope for an ADE protocol divided by the corresponding manual-dispense slope, with 95% confidence intervals (CI) and a reference line at 1.0 indicating parity with manual dispense. Red data points indicate significant deviation from manual pipetting (95% CI does not include 1). **h)** Lack-of-fit test for linearity across dispense volumes, larger values indicate stronger evidence that the fluorescence versus desired-volume relationship deviates from linearity. Black outlines, p < 0.05 by lack-of-fit test. **i)** Effect of evaporation on volume of H_2_O in source plates. **j)** Effect of moisture absorption on volume of anhydrous DMSO in source plates at 30% relative humidity.

### Sensitivity analysis of ADE using PickliPy.Assay

The applications below use both aqueous and DMSO stocks for dispensing from the same source plate. The manufacturer has strict recommendations for choosing the appropriate dispense protocol for the solvent. However, dispensing mismatched liquids with the same protocol may be more practical or faster than running the dispense twice with different protocols. Therefore, here we tested the magnitude of error introduced by using a mismatched protocol for solvent type. We used the concentration map feature of PickliPy.Assay to dispense increasing volumes of fluorescein solutions in H_2_O or in DMSO using ADE protocols for both aqueous and DMSO solutions (Fig. 2c-f). Mismatching aqueous and DMSO dispense protocols did not produce consistent volume deviations, although ADE dispensed less DMSO than manual pipetting (Fig. 2g). Importantly, the linearity of ADE was better in each case than that of manual pipetting (Fig. 2h) and we harness this in dose-response applications below.

Screening or experiments using live cells may follow a time course dictated by biology and, depending on the interval of dispense events, aqueous compound stocks may concentrate over time due to evaporation because of the high air flow in the dispense chamber of the Echo 650. This impacts experimental designs where the source plate remains in the instrument for a longer period for dispensing. Aqueous source volumes in 384PP plates evaporated at 3 µL/h during extended residence in the Echo chamber, regardless of door state (Fig. 2i). Conversely, anhydrous DMSO volumes increased by 0.75 µL/h (1.4%) at 30% ambient relative humidity (Fig. 2j). These effects can be mitigated by programmatically removing the plate from the acoustic dispenser and lidding in the wait periods (Fig. S4c).

These tests show that ADE can retain better dose linearity than manual pipetting even under pragmatic solvent-protocol compromises, while extended source-plate residence introduces predictable volume drift that should be managed in timed biological workflows. With these operating limits defined, we used PickliPy.Assay to move time-sensitive functional assays into pre-dispensed assay-ready formats.

### Application 1: Assay-ready ADE plates enable same-day multiplexed phenotyping of fresh human mitochondria

Using ADE, we exemplify how routine research lab assays can be enhanced by pre-dispensing customized assay-ready plates, sealing and freezing them for future use. The plates can be readily used at a time dictated by biology rather than by liquid handling instrument availability (Fig. 3). The preparation of assay-ready plates with acoustic dispensing has been reported in the past, indicating safe storage of nanoliter DMSO droplets at -18 °C ^14^. Here we demonstrate how PickliPy.Assay enables these workflows to allow uniform repeats and higher data density.

**Fig. 3.**
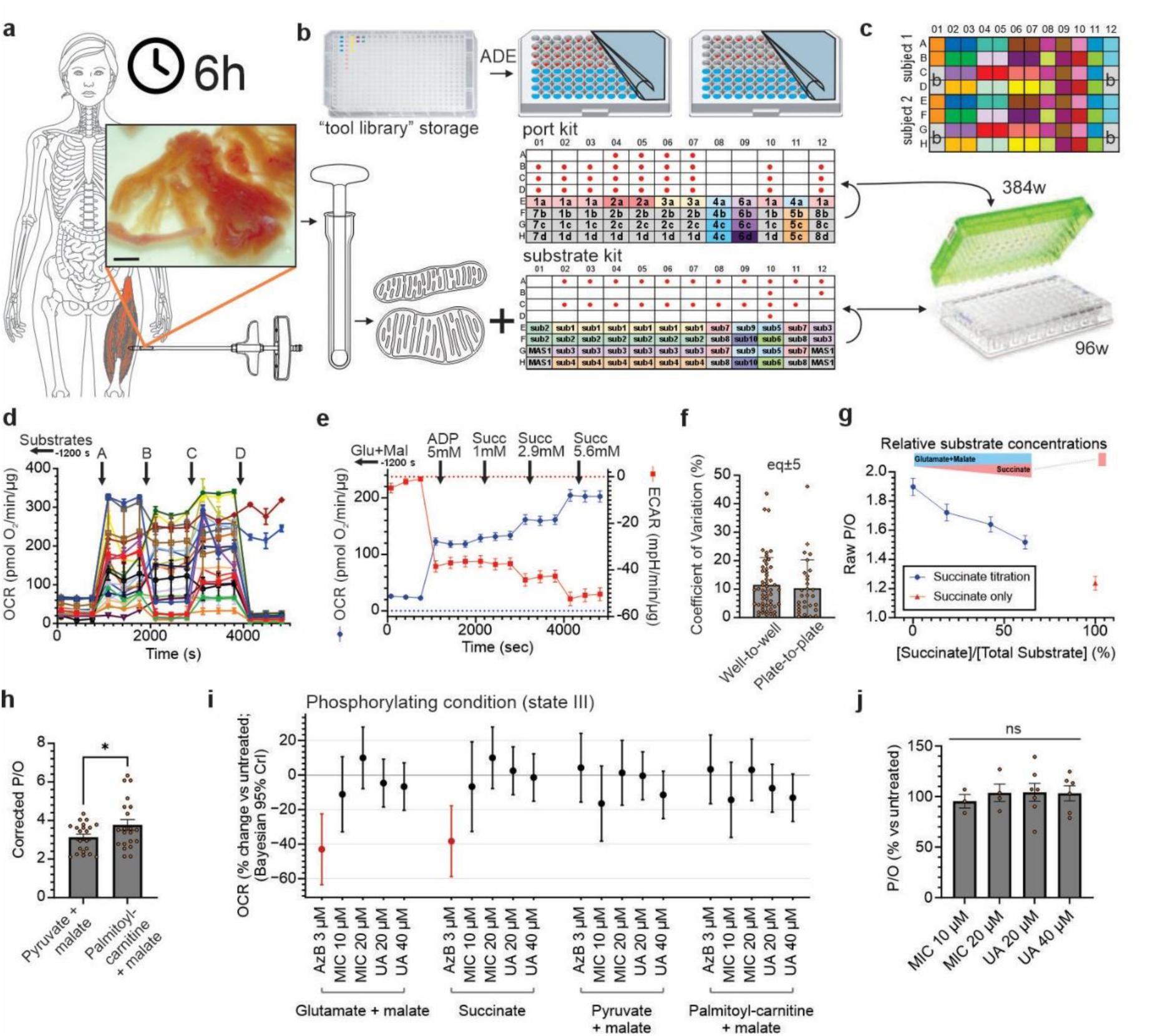
Assay-ready plates support time-sensitive, multiplexed human mitochondrial respirometry. **a)** Mitochondria isolated from human muscle biopsies (scale bar, 2 mm) from the vastus lateralis are useful for functional assays only for a short duration. **b)** Assay-ready plates were partly dispensed by ADE (red dots) and partly manually pipetted (blue and labeled wells). The “substrate kit” was added to the assay wells as the basal medium, while the “port kit” was added to the addition ports of the Agilent Seahorse XFe96 cartridge. **c)** The illustrated design uses two subjects per plate (top and bottom half) and 22 conditions with 2 well replicates and 4 background (b) wells for each. The XFe96 cartridge has 384 addition ports (four additions for each well). We used 8 distinct addition paradigms distributed across 12 columns and 4 rows (one for each addition) of the assay-ready 96-well plate **(b)**. Combinations of the 8 addition paradigms and 10 distinct substrate conditions made up the total 22 different conditions. **d)** Typical respirometry recording for a single subject (representative of 30 subjects) using the above 22 conditions (Wrobel et al., unpublished). A-D mark the four port additions that also correspond to four rows in the port plate in **(b)**. Data are mean ± SE, n = 2 wells in a representative plate. **e)** OCR and ECAR traces of a time course recording using the indicated port additions (Succ, succinate; total concentrations indicated), in mitochondria initially respiring on glutamate (5 mM) + malate (5 mM) added from the substrate kit plate about 20 min before the start of the baseline. **f)** Comparison of well-to-well (same ADE) and plate-to-plate (different ADE and XFe96 run) variations in OCR at a submaximal succinate concentration, so the reading is sensitive to the amount of succinate added as port addition. Data are mean ± SE for n = 30 subjects. eq±5, mean values were equivalent within a margin of ±5 percentage points (Welch TOST, α = 0.05; p = 0.0498). **g)** OCR and ECAR-derived P/O as a function of succinate content of substrate mix. Blue trace corresponds to data in **(e)**, the red symbol marks P/O calculated in phosphorylating respiration with succinate (5 mM; w/ rotenone, 2 µM) as substrate. **h)** OCR and ECAR-derived P/O in pyruvate (5 mM) + malate (1 mM) or palmitoyl-carnitine (15 µM ADE dispensed from DMSO stock) + malate (1 mM) substrate conditions. Values were corrected by a 1.31 multiplier calculated to match (g) to the theoretical values. *, p < 0.05 by paired t-test in n = 21 subjects. **i)** Effects of Azure B (AzB; 3.5 µM), MIC (10 µM, 20 µM) and UA (20 µM, 40 µM), on phosphorylating respiration in the presence of indicated substrates. Forest plot showing Bayesian posterior mean treatment effects with 95% credible intervals (CrI), grouped by substrate condition and adjusted for repeated subject measurements (typically 2 compounds), with FDR-significant effects shown in red. n = 5, 4, 5, 7, 6 subjects (one assay plate per subject) for the treatments in the indicated order. **j)** OCR and ECAR-derived P/O under the condition of palmitoyl-carnitine (12 µM), malate (0.78 mM), and pyruvate (0.49 mM) in the presence of UA (20 µM, 40 µM) and MIC (10 µM, 20 µM), expressed as % of the untreated from the same subject. No comparison was significant by ANOVA.

Due to the limited functional integrity of freshly isolated mitochondria, respirometry assays performed with the Agilent Seahorse Extracellular Flux Analyzer are extremely time sensitive and involve substantial liquid handling burden of 96 assay wells and 384 addition port wells (Fig. 3a-b). Routine runs of complex designs are therefore challenging and prone to manual errors if the layout is too complex. We demonstrate that ADE enabled complex bioenergetic phenotyping of human skeletal muscle mitochondria, where oxygen consumption rates (OCR) were surveyed in 22 different time course paradigms for each of the two subjects’ mitochondrial preparations per assay plate (Fig. 3c, d). In contrast, classical coupling (or respiratory control) and electron flow experiments were typically performed as one time course recording paradigm each^19^.

Experiments shown here took place within 6 h of the biopsies, including mitochondrial isolation and filling assay plates and cartridges. To accommodate this time frame, we designed and partly ADE-dispensed assay-ready “kit” plates (Fig. 3b). To benchmark this process, we examined a dose-response paradigm with the respiratory substrate succinate, one of the 22 traces in Fig. 3d, also shown as the average of 30 subjects in Fig. 3e (OCR trace). We calculated well-to-well and plate-to-plate variability from separate ADE runs at a submaximal succinate stimulation (Fig. 3f). The plate-to-plate variability (involving separate ADE) was equivalent to well-to-well variability.

The above approach of using ADE-prepared assay-ready kits converted Seahorse mitochondrial respirometry from a low-plex manual workflow into a reproducible, same-day multiplexed phenotyping platform. Below we harness this increase in experimental density by analyzing flux-derived metrics in a novel way.

### Measurement of P/O allows tracking substrate switching

The P/O ratio describes the stoichiometry of OXPHOS; the number of ATP molecules synthesized per consumed oxygen atom. Its value may be used for tracking substrate oxidation pathways in isolated mitochondria^20^. It is a classical respirometry metric based on tracking the phosphorylation of a finite amount of added ADP, but a high-throughput assay has been missing because of the difficulty of tracking such a finite challenge in microplate-based respirometry.

Therefore, here we applied a novel, flux-based approach to calculating P/O ratio in isolated mitochondria from coupled OCR and extracellular acidification rate (ECAR) data using a Seahorse XF analyzer. This calculation is possible because phosphorylation of ADP results in net alkalinization. To demonstrate this, we used the above paradigm with incremental ADE-dispensed succinate additions, and captured substrate switching from complex I-driven glutamate + malate respiration to complex II-driven succinate oxidation (Fig. 3e, g). As succinate increasingly drove respiration, P/O declined from the higher complex I value (theoretical maximum of 2.46^20^) toward the lower complex II value (1.64^20^; Fig. 3g and Table S1). A deviation from the theoretical maximum is likely due to a known limitation imposed by the Seahorse XF analyzer’s absolute scaling^21,22^ and therefore we show corrected values below using the empirical factor of 1.31 ± 0.03 (mean ± SE, n=30) that was calculated by scaling the measured values in Fig. 3g to their corresponding theoretical values. Pyruvate + malate and palmitoyl-carnitine + malate yielded significantly different P/O values, allowing tracking of substrate switching (Fig. 3h). The alkalinization-derived P/O, accounting for TCA-cycle-related acid formation was higher than expected (Table S1), consistent with prior studies^23^. The increased throughput provided by our method can be harnessed in future mechanistic studies to understand the higher-than-expected P/O. To test high-density respirometry and microplate-scale P/O readouts in practice, we analyzed the three small molecules relevant to aging interventions in this context.

### No direct, acute bioenergetic effects for Urolithin A and mitophagy-inducing coumarin (MIC)

Urolithin A (UA) has been studied extensively as a mitochondrial quality-control enhancer, with measurable effects on mitochondrial biomarkers and improved musculoskeletal outcomes in humans after chronic treatment^24,25^. While the best-supported mechanism of action for UA is increased mitophagy through activation of the PINK-1 pathway^26^, direct experiments in isolated human mitochondria have not, to our knowledge, been reported. Using the above methodology, we tested the direct, acute effects of UA and the recently described mitophagy-inducing coumarin (MIC)^27^ in human muscle mitochondria from 30 human subjects (Fig. 3i). To this end, we used PickliPy.Assay to uniformly dispense 30 nL of MIC, UA, and Azure B (a redox-active methylene blue derivative as a positive control^28^) stocks into additional storage plates and manually transferred “substrate kit” plate media through these wells before filling the assay plate. After a 30 min pretreatment, neither UA (20, 40 µM) nor MIC (10, 20 µM) affected Complex I- or II-driven respiration in human skeletal muscle mitochondria. In contrast, Azure B (3.5 µM) strongly inhibited pure Complex I or II-mediated respiration, but not respiration with substrates malate + pyruvate present. In the latter substrate combination, TCA-cycle reactions that generate NADPH may fuel antioxidant defenses to counter the pro-oxidant effects of Azure B. UA has been reported after 24 h treatment to increase fatty-acid-oxidation-driven OCR in differentiated C2C12 myotubes^26^, and to shift gene-expression signatures related to β-oxidation^29^. Thus, these effects required longer-term incubations in intact cells, and consistently, we found no acute direct effects of UA on isolated human skeletal-muscle mitochondria, including the P/O ratio of mixed substrates (pyruvate + palmitoyl-carnitine; Fig. 3j). Having established ADE-prepared Seahorse assays for isolated mitochondria, we next broadened PickliPy’s applications towards live-cell time-lapse fluorescence microscopy.

### Application 2: ADE-prepared reservoir plates extend live-cell membrane-potential assays

Human dispersed pancreatic islet cells from post-mortem donors have a limited time window (∼10 days) of primary culturing for functional assays^30^. Here we aimed to maximize the number of assay conditions by microplate-based multiplexing (Fig. 4a) during fluorescence time-lapse microscopy of mitochondrial and plasma membrane potentials (ΔψM and ΔψP)^31,32^. To challenge cell cultures with glucose and added bioenergetic or metabolic modulators, we previously designed a partial-96-well microplate paradigm with multiple manual, multi-channel pipetting steps during time courses on a standalone fluorescence microscope^32^. To minimize added vehicle, we used an Excel-based calculation table^32^ to prepare reservoir microplates (Fig. 4c) holding excess amounts of 2×-concentrated treatments and we manually applied this to the assay plate during time course recording (Fig. 4d). Manual preparation of these reservoir plates was time-consuming and challenging, involving pipetting small volumes with limited accuracy. Here we describe a modification of the preparation of these reservoir plates using ADE-dispensed assay-ready plates (Fig. 4b). Carrying out the live-cell time-lapse experiments remained manual. To this end, we created PickliPy.BlueTable, a specialized version of PickliPy.Assay to generate assay-ready plates based on the above calculation worksheet.

**Fig. 4.**
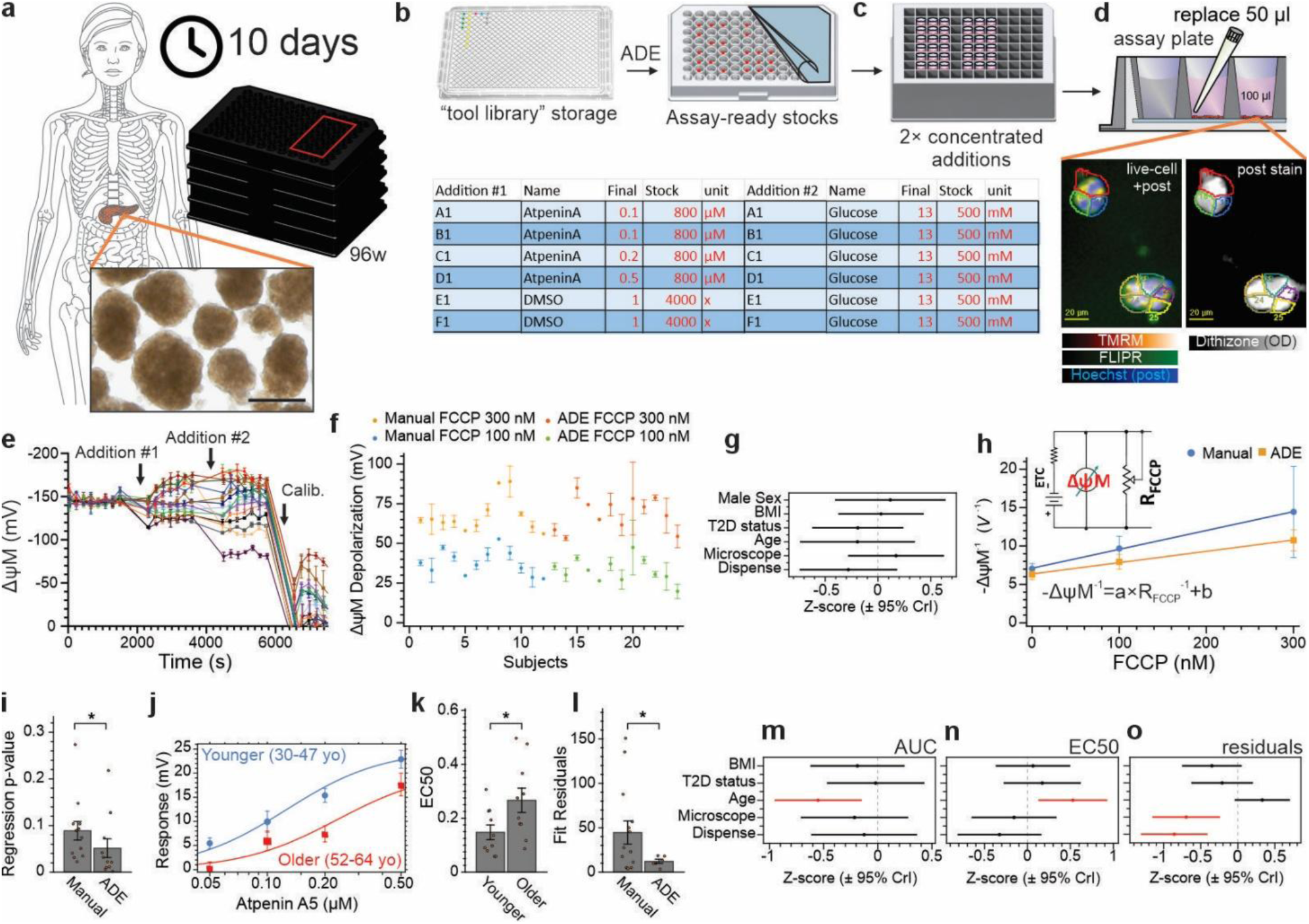
Assay-ready plates support manual liquid handling in microplates during fluorescence microscopy time course recordings. **a)** Human primary islet cell cultures have a limited time window for assaying. Image shows intact islets before dispersion, scale bar, 200 µm. **b)** Assay-ready plates were dispensed with ADE, using PickliPy.BlueTable, and then dissolved in assay medium before transferring to the **c)** assay reservoir. **d)** Fluorescence microscopy was performed In the cultures with manual half-replacement of well content for challenges and modulations. Micrographs: glucose-stimulated β-cells during a live-cell time course with TMRM and FLIPR staining, overlaid by Hoechst 33342 fluorescence captured post time lapse. β-cell identity was defined by dithizone staining (staining insulin-bound zinc) post time lapse on the same cells, shown as an optical density (OD) image. **e)** Representative 18-condition time-course recording (of 91 experiments using ADE). Data are mean ± SE of n = 2 field-of-view replicates per well. **f)** ΔψM depolarization triggered by 100 nM and 300 nM FCCP (w/o BSA or FBS) in pancreatic β-cells, in the presence of 16 mM glucose, showing data from 24 subjects that were recorded independently. Samples from subjects 1-12 were assayed with manual dilution of FCCP, whereas subjects 13-24 were assayed using ADE with the workflow illustrated in **(b-c)**. Data points are mean (± SE), n = 1-3 of experimental replicates per subject. **g)** Bayesian multivariate linear regression analyzing factors contributing to the magnitude of ΔψM depolarization by 100 nM FCCP. **h)** Regression analysis per dispense modality on the relationship of -ΔψM^-1^ and FCCP concentration ∼ R_FCCP_^-1^. Data points are mean ± SE for all subjects per dispense modality, n = 12 and 12. Inset: simple electric model of mitochondria. Ohm’s law dictates a linear relationship between R_FCCP_^-1^ and -ΔψM^-1^. **i)** Analysis of p-values of per subject fits corresponding to the average fits shown in h. *, p < 0.05 by Kruskal-Wallis test. **j)** Dose-response curves of ΔψM depolarization triggered by the SDH inhibitor Atpenin A5 in the presence of 16 mM glucose in pancreatic β-cells from younger (30-47 yo; n = 10) and older (52-64 yo; n = 10) subjects. Data points are mean ± SE; average fit curves shown. **k)** Comparison of EC_50_ values of individual fits between younger and older subjects, corresponding to **j)**. Data points are subjects. **l)** Comparison of fit residuals between experiments performed with manual and ADE dispensing. **k,l)** *, p < 0.05 as compared by Welch test. **m**-**o)** Bayesian multivariate linear regression analyzing factors contributing to variations in metrics of per subject fits in **(j)**: **m)** area under the curve (AUC); **n)** EC_50_ values (corresponding to **k**); **o)** fit residuals (corresponding to **l**). Each CrI measures effect corrected for all other covariates. Red, significance determined by 95% CrI excluding 0, n = 20 subjects.

The ADE-generated reservoir plates were used in 18-well, two-addition time-course recordings of ΔψM in human pancreatic β-cells (Fig. 4e). We tested the repeatability of delivering bioenergetic modulators in a cohort study by analyzing the depolarization of ΔψM by submaximal 100 and 300 nM concentrations of the uncoupler FCCP (Fig. 4f; one assay condition from experiments like the one shown in Fig. 4e). Data in Fig. 4f were collected over 10 years, with the first half using manual and the second half using ADE dispensing. The manual method involved preparation of diluted stocks for each concentration, while ADE dispensed a single concentrated compound stock backfilled with DMSO (total of 100 nL). Although effect size varied across subjects, it was not associated with the mode of dispense (manual vs ADE) after controlling for clinical covariates and microscope setup (Fig. 4g). To benchmark dose-response dispensing, -1/ΔψM was graphed as a function of FCCP concentration. Based on Ohm’s law, a linear relationship is expected (Fig. 4h). In this historical, sequential comparison, ADE-dispensed FCCP dose responses were closer to the expected linear relationship than earlier manual-dispense responses (Fig. 4i), although dispense method was confounded with experimental period. This technical gain in micro-scale dose-responses was further harnessed after asking how human donor biology alters ΔψM regulation.

### ADE improves dose-response precision and reveals age-associated kinetic changes in human β-cells

Succinate dehydrogenase (SDH) is a component of the electron transport chain (ETC) that is thermodynamically capable of generating higher proton motive force than Complex I^33^, and its activity is required for insulin secretion^34^. Age-related changes in mitochondrial respiratory function and complex-II-linked phenotypes appear tissue-dependent, with declines reported in human liver mitochondria ^35^, conflicting findings in muscle due to the impact of exercise^36^ and no change in heart^37^. Using the above approach, we recorded 4-point dose-response curves of mitochondrial depolarization by SDH-inhibitor Atpenin A5 in pancreatic β-cells, seeking to interrogate kinetic differences between subjects spanning ages 30-64 years (Fig. 4j). Older donor age was associated with a smaller Atpenin A5 dose-response area under the curve (AUC), consistent with a reduced SDH-inhibitor depolarizing effect (Fig. 4m). Type 2 diabetes (T2D) status and technical variables, such as manual versus ADE dispense were not associated with the outcome. This difference was due to increased EC_50_ of Atpenin A5 (Fig. 4k, n). To benchmark the dispense, we compared Hill equation fit residuals between manual and ADE dispenses. ADE significantly improved the fits (Fig. 4l, o), suggesting better accuracy than manual dispense of intermediate dilutions. In summary, dose-response analysis of Atpenin A5 revealed that ΔψM in older human donor β-cells is less depolarized by SDH inhibition, suggesting an age-associated change in SDH kinetics or ΔψM control. After these focused assay-ready workflows, we next tested PickliPy.Screen in an exploratory mitochondrial phenotyping workflow.

### Application 3: Three-dose mitochondrial morpho-functional screening with library reformatting

High-content mitochondrial phenotyping has become a productive strategy for compound profiling and mechanism discovery. Recent studies have combined mitochondrial morphology, membrane potential, pH-dependent mitophagy reporters, and morphomic feature extraction to classify mitochondrial perturbations, infer drug mechanisms, or identify mitochondrial modulators ^38–40^. Building on these studies, we used mitochondrial phenotyping to benchmark assay-window robustness across functional and morphological endpoints within an end-to-end workflow spanning ADE-based library handling, live-cell imaging, and downstream analysis.

To combine mitochondrial morphology with a live functional readout, we generated HEK293 cells that stably and uniformly expressed mitochondrially targeted low-affinity ATP sensor AT1.03, mito-ATeam (Fig. 5a-c). Using a wash-free paradigm for drugging, culturing, and fluorescence microscopy imaging of live cells, we implemented a screen for broad mitochondrial bioenergetic and morphological effects of small molecules. This may be used as a pre-screen to select compound concentrations, identify mitochondrial liabilities, and define compound subsets for more complex follow-up screens using PickliPy, or as we show below, serve as a sensitive mitochondrial phenotyping platform. Library compounds were reformatted from three source plates into 13 destination assay plates at three final concentrations per compound (1, 3, and 10 µM), with positive and vehicle controls added to each plate (Fig. 5d, e). This was controlled by a single picklist. After 48 h of culturing, imaging was performed using scheduled robotic loading of microplates with live cells from the TC incubator to the environment-controlled recording chamber of the microscope (Fig. 5f and Fig. S5a-c). Next, we stress-tested this design-to-picklist coordinated workflow by analyzing controls included in the screen.

**Fig. 5.**
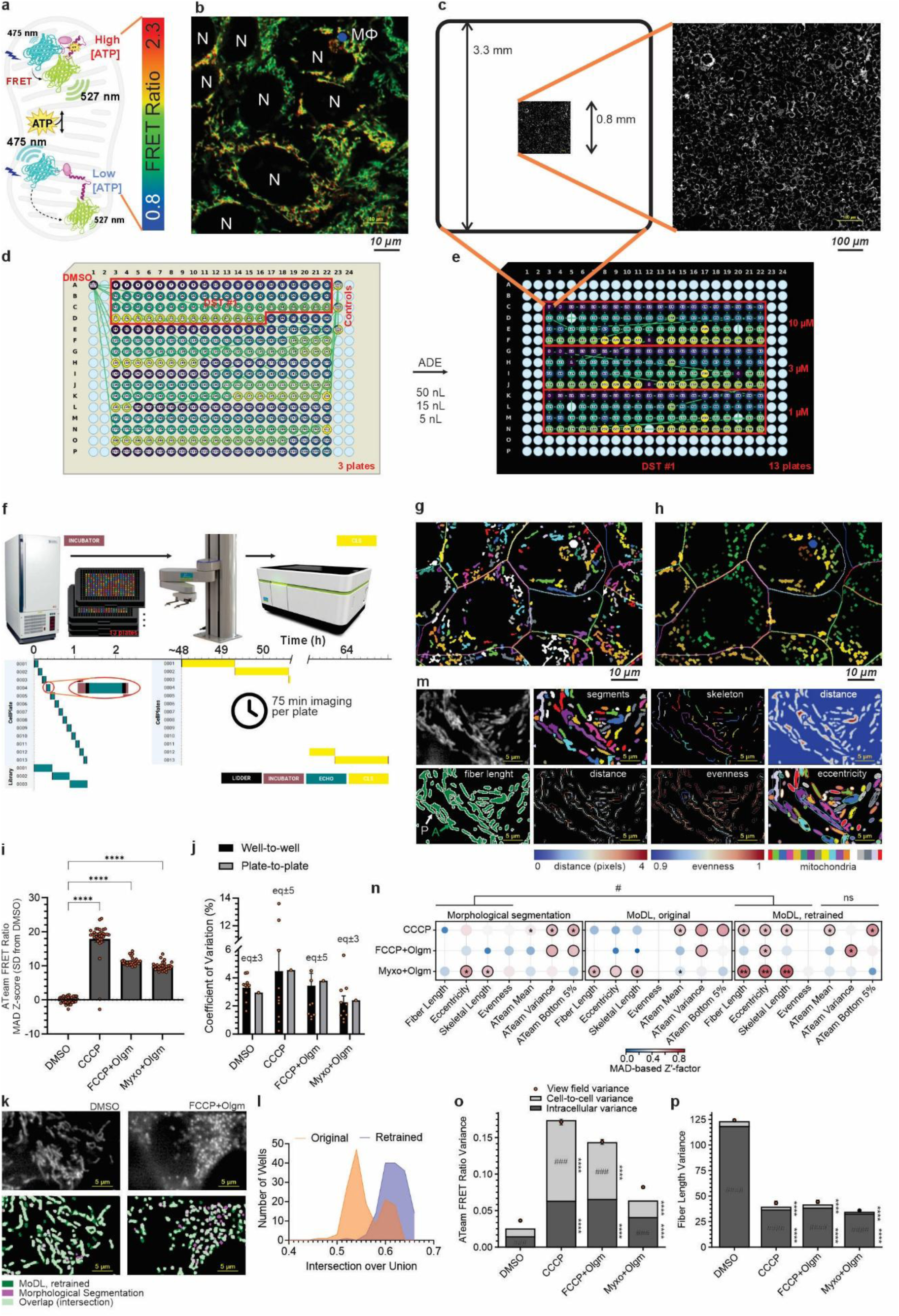
Small-molecule library screening with reformatting and three doses per plate for modulators of mitochondrial [ATP] and morphology using PickliPy.Screen. **a)** mito-ATeam senses mitochondrial ATP levels in the matrix. **b)** Typical confluent culture of HEK-mAT1.03 cells used for screening, shown as an intensity-modulated fluorescence ratio image. N, nucleus; MΦ, mitophagy. **c)** Stitched (4×4 tiles) typical view field of mito-ATeam intensity showing uniform expression, visualized proportionally to the well size of the 384-well plate. **d)** Source and **e)** destination plate movements during ADE, showing the first library and first cell culture plate (DST #1). Red frame indicates library compound wells dispensed into DST #1, and the three doses used in the destination. **f)** Timing of dosing and plating. Plates were moved by the PFlex robot between the incubator and the Echo 650 or the Operetta CLS for maintained environment control. **g)** Analysis of mitochondrial morphology (colored segments; using morphological segmentation) and cell shapes (colored outlines). Representative image, corresponding to **(b)**. **h)** Analysis of intracellular heterogeneity of matrix [ATP] by calculating the ratio of fluorescence intensities over visually continuous mitochondria. Pseudo color coding, as defined in **(a)**. **i)** Matrix [ATP] in positive controls normalized to vehicle-treated wells. Data points are wells pooled from all plates. ****, p < 0.0001 by Kruskal-Wallis multiple comparison. **j)** Comparison of well-to-well and plate-to-plate coefficients of variation (CoV) of matrix [ATP] in control wells. Data points are plates, or single data points are shown for plate-to-plate variations. eq±3 and eq±5, mean values were equivalent within a margin of ±3 and ±5 percentage points, respectively (Welch TOST, α = 0.05; p < 0.05 for n=10 plates) **k)** mito-ATeam fluorescence images (top) and overlay (bottom) of MoDL segmentation with retrained network weights (green; see results with original weights in Fig. S5e) and morphological segmentations (purple) in the indicated conditions. **l)** MoDL segmentation with in-dataset retrained network weights provided more consistent overlaps with the unbiased morphological segmentation results in pooled conditions than using the pretrained weights, as indicated by the unimodal distribution (blue) of the intersection over union values, calculated as shown in **(k)**. **m)** Mitochondrial morphomic analysis using MoDL segmentation. Top row from left: mito-ATeam fluorescence; segmented image obtained by retrained MoDL with watershed segmentation of the probability map; skeletonized segments; distance image, where pixel values are calculated as the distance to the nearest background pixel. Bottom row from left: fiber length is calculated from the area (A) and perimeter (P) of segments; distance image masked with skeleton (overlay: perimeter of segments); mitochondrial skeletons colored according to the normalized Shannon effective-number evenness values (higher values indicate more uniform thickness); eccentricity was calculated from the best-fit ellipses. **n)** Robust MAD- (median absolute deviation) based Z’-factors (*Z*′_MAD_) of screening with metrics illustrated in **(m)**, comparing the positive controls to DMSO, for three image analysis pipelines from left: morphological segmentation, MoDL with original network weights, and MoDL with in-dataset retrained network weights. All MoDL analysis used post-processing of probability maps, except in the corresponding analysis in Fig. S5f. Dots indicate *Z*′_MAD_ > 0; black outlines, *Z*′_MAD_ > 0.5, * and **, 95% CI of *Z*′_MAD_ above 0.3 and 0.5, respectively; #, significant improvement comparing retrained MoDL to morphological segmentation, using Holm-adjusted one-sided bootstrap tail probabilities < 0.05 for morphology or ATeam metric classes; ns, not significant. **o-p)** Variance hierarchy decomposition to distinguish contribution of cell-to-cell and intracellular variation (stacked bars) to the total, field-of-view-level variance (data points) of **(o)** ATeam FRET ratio and **(p)** Mitochondrial fiber length (see also Fig. S5g for eccentricity). Data are from n = 56, 30, 30, 31 wells pooled from 13 microplates for DMSO (1:1000 v/v), FCCP + Olgm (oligomycin; 4 µM + 2 µg/mL), Myxo + Olgm (4 µM + 2 µg/mL), CCCP (10 µM), after quality control excluding wells with < 100 cells. ****, BH FDR q ≤ 0.0001 compared to the same metric in DMSO. ### and #### indicate the larger component at BH FDR q ≤ 0.001 and 0.0001, respectively, within each stack using bootstrapping.

### Mitochondrial matrix [ATP] sensitively tracks bioenergetic modulations

Mitochondrial matrix [ATP] was quantified after cell and mitochondrial segmentation, restricting analysis to mitochondrial regions within in-focus cells (Fig. 5g, h). As strong bioenergetic perturbation controls, we used the uncoupler CCCP alone, and either FCCP or the complex III inhibitor myxothiazol in combination with the ATP synthase inhibitor oligomycin to prevent mitochondrial hydrolysis of glycolytic ATP. These controls were dispensed from wells included on each library source plate (Fig. 5d). These strong respiratory perturbation controls produced highly significant shifts in DMSO-normalized well-level mito-ATeam FRET ratios (Fig. 5i), but seemingly counterintuitively, increased matrix [ATP] concentration. This is expected because, matrix ATP concentration is not a direct proxy for oxidative ATP synthesis. Instead, matrix adenine nucleotide ratios are shaped by the electrogenic adenine nucleotide translocase (ANT), which normally supports ATP export and ADP import and thereby contributes to a lower matrix ATP/ADP ratio than in the cytosol. In highly glycolytic HEK293 cells, this behavior is consistent with reversal of ANT under mitochondrial depolarization, allowing cytosolic ATP import into the matrix in exchange for ADP^41^, and with evidence that glycolytically produced ATP can support matrix ATP during respiratory inhibition^42^.

To benchmark ADE, we analyzed plate-to-plate variations that were not greater than within-plate well-to-well variations (Fig. 5j), indicating that the scheduled timing and environmental control limited systematic plate-position and plate-order effects. Positive-control responses were detected as expected in 90 wells; however, two to three CCCP control wells phenotypically failed for unknown reasons with no logged dispense error.

Thus, mito-ATeam detected expected bioenergetic perturbations across the screen with plate-order effects no larger than within-plate variation, supporting matrix ATP as a sensitive and practical functional endpoint. Because this endpoint depends on accurate mitochondrial segmentation, next we used the above data to optimize related image analysis.

### Pretrained deep-learning segmentation can be biased by screen-induced biology and improved by in-dataset retraining

Deep learning has improved mitochondrial segmentation in fluorescence microscopy imaging and has recently enabled prediction of mitochondrial functional states from morphology^43^.

MitoSegNet was trained on mixed mitochondrial phenotypes^44^, and MoDL extended and updated this approach^43^. Importantly, pretrained models encode the cell type, probe behavior, resolution, annotation style, and phenotype distribution of their training data. Their performance in HTS must therefore be tested against the specific biology generated by the screen.

Here we evaluated the performance of two segmentation approaches on the above data: a spatial filtering plus morphological segmentation-based method that we previously designed for quantitative measurements outlining mitochondria at their half-maximal intensity^31^ and the U-RNet+ architecture MoDL^43^ that reflects the mitochondria-outlining segmentation style of 15 biologists using healthy cells. We used the vehicle and positive-control conditions to compare these analysis approaches (Fig. 5k and Fig. S5d).

The positive controls induced severe mitochondrial damage with profuse fragmentation. Under these conditions, the conventional morphological segmentation and the pretrained MoDL model diverged: the pretrained network missed many punctate mitochondria generated by the noxious controls (Fig. S5d arrows). This was reflected in a bimodal distribution of intersection-over-union values between the two segmentation outputs across conditions (Fig. 5l). We therefore retrained the original MoDL weights using 445 vehicle and positive-control image patches, with algorithmically generated and visually quality-controlled ground truth masks (Fig. S5e). Dataset-specific retraining improved mask overlap between the morphological-segmentation and deep learning approaches across both control and damaged states (Fig. 5l), demonstrating that biologically induced out-of-distribution morphology can bias segmentation and that this bias can be mitigated by modest in-assay retraining.

To determine whether the improvement in image segmentation translated into better screening performance, we benchmarked four endpoint classes based on distinct measurement principles: perimeter- and area-derived fiber length, PCA-based eccentricity, a skeletonization-derived mitochondrial length and thickness evenness (Fig. 5m), and mito-ATeam FRET-ratio statistics measured over segmented mitochondrial regions (Fig. 5h). We found a significant improvement when MoDL was retrained on in-dataset controls, compared with conventional morphological segmentation (Fig. 5n and Fig. S5f).

### Mitochondrial matrix [ATP] varies cell-to-cell, whereas morphology is dominated by intracellular heterogeneity

The retrained workflow analyzed 5.3 million mitochondrial profiles from 92,000 cells across 147 wells in a homogeneous HEK293 culture. We used hierarchical variance decomposition to determine whether variability arose mainly between cells or among mitochondrial profiles within cells. Mito-ATeam FRET-ratio variance contained both cell-to-cell and intracellular components (Fig. 5o), whereas fiber length and eccentricity variance were explained predominantly by intracellular heterogeneity (Fig. 5p and Fig. S5g). Noxious bioenergetic perturbations increased heterogeneity in matrix [ATP] but made mitochondrial morphology more uniform, consistent with collapse of a normally heterogeneous mitochondrial network into a more uniformly damaged, fragmented state.

These results and related benchmarks in Fig. 5n indicate that heterogeneity itself can serve as a screening endpoint, beyond changes in population-average intensity or morphology. The observed cell-to-cell variability in matrix [ATP] further argues that mitochondrial high-content screens should optimize image acquisition for the number of sampled cells and fields of view, rather than rely on small view fields that may undersample heterogeneous cellular responses, even in a uniform cell line such as HEK293. The following and last application harnesses cell-to-cell heterogeneity and precise ADE dosing to characterize compound effects on mitochondrial biology.

### Application 4: ADE-dispensed dose responses reveal dose-range-specific mitochondrial morpho-functional effects

To demonstrate a follow-up for the screen, we created a cherry-picked daughter plate for a subset of the original library plus controls, including one intermediate dilution of compounds on the same plate using PickliPy.Screen (Fig. 6a). Then, in a separate workflow, we used a nine-point dose-response layout with constant DMSO concentration and randomized repeats as shown in Fig. 1e-h. Using the assay illustrated in Fig. 5f-h, m, we analyzed a set of compounds that we selected from the original screen based on the strong phenotypes they induced.

**Fig. 6.**
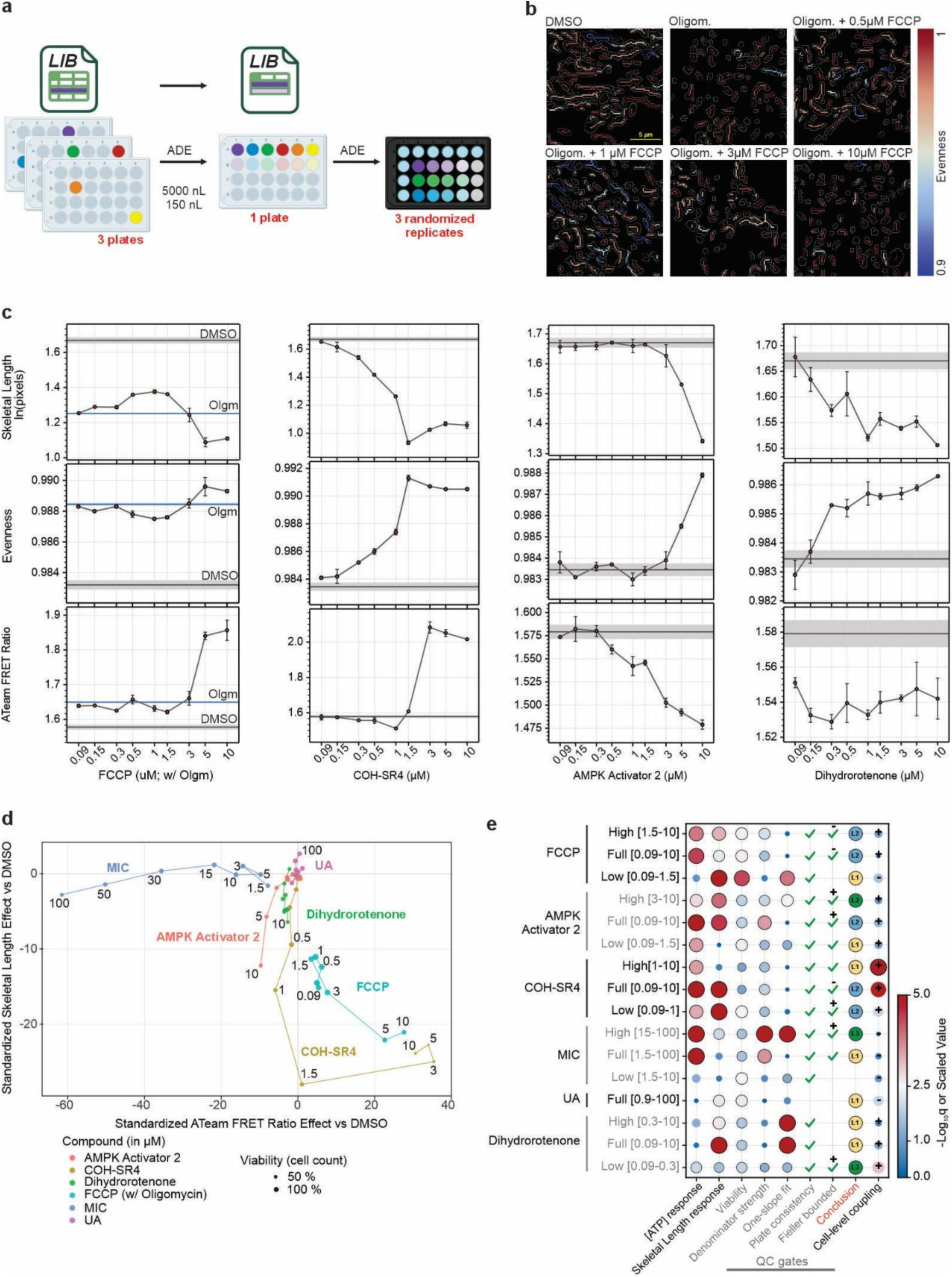
Dose-response experiments using PickliPy.Screen reveal morpho-functional coupling in HEK293 cell mitochondria. **a)** Using a shortlisted LIB table, compounds of interest were ADE-dispensed from the original library plates into one daughter plate at two volumes allowing for one intermediate dilution (DMSO was manually added). Then, using a modified LIB table dose-responses were dispensed as shown in Fig. 1e-h and the mito-ATeam HEK293 cells were cultured for 48 h. **b)** Processed images showing mitochondrial skeletons colored according to mitochondrial evenness values. Outlines indicate the perimeter of mitochondrial segments. Frames correspond to data points in FCCP dose-response in the presence of oligomycin below. **c)** Dose-response relationships showing mean ± SE of n = 3 plate replicates with randomized layouts. FCCP titration in the presence of oligomycin (2 µg/mL) and 10% FBS. Gray shaded areas show the DMSO control (median ± MAD n = 66), and for FCCP titration the DMSO + Olgm (n = 6) control is shown in blue. COH-SR4 (no oligomycin) behaved similarly to the uncoupler FCCP, but with a low-dose mitochondrial fragmenting effect. AMPK Activator 2 demonstrated a low-dose ATP-lowering effect with morphology impacted more at higher doses. In contrast, dihydrorotenone, a complex I inhibitor, resulted in only modest ATP lowering and mitochondrial shortening. **d)** Function-morphology effect-space trajectories. Each point shows the paired well-averaged effect, standardized relative to plate-matched DMSO controls. Lines connect increasing concentrations, providing a dose-dependent path through the function-morphology space. Distinct trajectories indicate that mitochondrial length did not simply mirror the ATeam FRET ratio across perturbations; instead, the relationship between functional and morphological phenotypes varied by compound and dose range. **e)** Dose-response co-response hierarchy for ATeam FRET ratio and skeletal length. Rows summarize full-range and optimized breakpoint-defined high and low concentration windows for each compound. Columns show the evidence used to classify morpho-functional coupling. Black outlines mark accepted gates (FDR q < 0.05 for responses, for others see Methods). Binary endpoints are marked by checkmarks. One-slope adequacy, whether a single straight slope adequately fits; Plate consistency, slope-sign agreement in at least two of three plates; Fieller bounded, slope-ratio confidence interval with sign shown if the interval is completely above/below zero. A Fieller interval for the morphology/[ATP] local-slope ratio entirely above zero (indicated by a +) shows that a dose-dependently decreasing matrix [ATP] was associated with shortening mitochondria. The Conclusion column reports the hierarchy level: L1, single-endpoint or dissociated response; L2, dose-associated co-response; and L3, proportional local coupling. Cell-level coupling was defined as the effect of the compound on the within-well association between the functional readout and the morphological endpoint, with direction indicating whether the coefficient of this correlation increased or decreased by the drug. Responses and coupling are provided in -log_10_q values capped at 5 (see other morphology metrics in Fig. S7).

### Uncoupling of OXPHOS produces biphasic functional and morphological shifts

The primary screen indicated distinct dose-response characteristics among the library compounds, including biphasic response to uncouplers (Fig. S6a). Therefore, we evaluated the nine-point dose response for the uncoupler FCCP (in the constant presence of oligomycin), the uncoupler and AMPK-activator COH-SR4^45^ and the biguanide AMPK Activator 2 (compound 7a in ^46^). FCCP at low concentrations partially reversed the mitochondria-fragmenting effects of oligomycin and lowered matrix ATP levels (Fig. 6b-c). However, above 1.5 µM, likely at a point where ΔψM collapses, mitochondria became more fragmented and matrix ATP increased. This is consistent with a differential sensitivity of ATP-synthesis and ANT to mitochondrial depolarization^41^. COH-SR4, with known uncoupler properties behaved similarly at high concentrations. MIC at 48 h dose-dependently decreased mitochondrial FRET (Fig. S6b), despite the lack of acute, direct mitochondrial effects (Fig. 3i), and UA had no consistent effect up to 30 µM and slightly elongated mitochondria at 50-100 µM. ATeam is pH sensitive, thus some of the observed larger decreases in ATeam FRET ratio may be due to upregulated glycolysis and medium acidification during the 48 h treatment.

Each of the observed dose-response series (Fig. 6c and S6b) showed complex, divergent effects on ATP levels and morphology and motivated an explicit analysis of whether morphology and function are quantitatively coupled below.

### Dose-range-specific morpho-functional coupling

How mitochondrial function relates to mitochondrial morphology is a long-standing question. Here, precise ADE dose-response recordings allowed us to organize paired functional and morphological phenotypes from high-content imaging into a response hierarchy that makes structure-function claims explicit. The hierarchy separates three increasingly stringent levels based on well-averaged responses to escalating drug concentrations: **L1**, a single-endpoint or dissociated response, in which only one metric moves; **L2**, dose-associated co-response; and **L3**, proportional coupling that allows prediction of one metric from the other. The dose-response path through function-morphology space followed distinct trajectories, indicating that the structure-function relationship varied by perturbation and dose range (Fig. 6d), and the hierarchy analysis made this quantitative (Fig. 6e).

A dose-dependently decreasing matrix [ATP] was associated with shortening mitochondria for AMPK Activator 2 at 3-10 µM, MIC at 15-100 µM, and dihydrorotenone at 0.09-0.3 µM (L3; Fieller +; Fig. 6e). Effects of FCCP at the high range of 1.5-10 µM, where bioenergetic collapse is expected, showed morpho-functional association, but were non-linear when measuring mitochondrial length (L2). However, matrix [ATP] was proportionally coupled in this range with thickness evenness (L3; Fig. S7). Interestingly, the nonlinear trajectories of FCCP and the other known uncoupler, COH-SR4, included dose ranges in which increases in matrix [ATP] coincided with mitochondrial shortening. At the cell level, DMSO wells showed a positive within-well association between ATeam FRET ratio and skeletal length (well-clustered BH q = 7.7×10^-9^ from 66 DMSO wells). Thus, cells with relatively higher matrix [ATP] had longer mitochondria. COH-SR4 significantly strengthened this association (Fig. 6e), that may be explained by increased cell-to-cell heterogeneity (Fig. S7c).

Altogether, ADE provided precise dispensing and enabled complex randomization of dose-response layouts unpenalized by human error. These collectively supported a dose-resolved morpho-functional analysis, indicating that mitochondrial elongation was associated with higher matrix [ATP], except for the case of bioenergetic collapse in the presence of high concentrations of uncouplers.

### PickliPy supports agent-assisted experimental design and troubleshooting

To shorten user learning curve providing an interactive troubleshooting, or to further the complexity of experimental designs, such as using design of experiment (DOE) principles, we created an agentic skill with the ability to make design files, downstream picklists and metadata using PickliPy (Fig. S8).

## Discussion

Combining ADE with the flexible control provided by PickliPy has changed how we design and execute both bench-scale experiments and high-throughput screens. The largest gains at the bench came from using assay-ready plates to treat cell cultures and mitochondria in microplates, which increased data density and improved replicate uniformity. Many routine assays with live samples are time-sensitive and rely on specialized instruments that are not integrated into automated workflows. By contrast, integrated robotic workcells are often housed in core facilities, operated by specialist staff and are less easily scheduled for the precise start times required by complex assays. Assay-ready plates bridge these two workflows: they enable centralized, programmable preparation while preserving the flexibility of bench-side execution. As shown here, one important application is routine assays in cohort studies, where customized assay-ready plates can multiplex far more conditions than would be practical by hand pipetting.

For high-throughput screening, we focused on providing assay design tools in the absence of an information-management backend. We described a low consumables cost, wash-free screen that may precede a costlier and more laborious immunocytochemical detection of specialized biology using a narrowed-down chemical library at optimized concentrations for each compound, harnessing the shortlisting and dosing capabilities of PickliPy.Screen. Our approach extends previously demonstrated^43,47,48^ morphological phenotype space analysis with functional aspects, and here we demonstrate that modern AI tools and complex downstream analysis can be done at scale.

A major theme here was increasing the depth of bioenergetic analysis at scale in cohort studies and screening contexts. To date, a handful of candidate small-molecule mitochondrial interventions have been identified, with aspirations for mitigating disease or age-related decline, including UA^24^ and MIC^27^. Given the typical ambiguity of the mechanism of these interventions, immediate applications of our bioenergetics toolset are to understand existing mitochondrial enhancers and screen for novel ones. This requires a broader set of metrics for evaluating mitochondrial function. One of these is heterogeneity of observed variables, such as matrix [ATP], at the cellular and subcellular level. Metabolic stress tends to increase mitochondrial heterogeneity. However, heterogeneity is not necessarily detrimental, cells can maintain metabolically distinct mitochondrial subpopulations for different purposes^49^.

Broadening the within-cell mito-ATeam distribution after drug treatment therefore may reflect the emergence of stressed or compensating mitochondrial subpopulations.

An important theme of our approaches was using precise dose-response assays that were more accurately performed in the small assay volumes with ADE than with manual pipetting. This enabled us to design and apply kinetic analysis in isolated mitochondrial and cellular systems and a dose-response co-response hierarchy analysis in the high-throughput screening context associating mitochondrial morpho-functional metrics, providing a novel toolset to seek mechanistic understanding of mitochondrial biology.

In summary, the presented experimental design framework (PickliPy) provides a modular and accessible solution to a common bottleneck in ADE-based workflows and supports broader efforts to improve reproducibility and data integrity in bench-scale and high-throughput screening environments. The resulting increase in data density, combined with modern data analysis tools, enabled a more sensitive and comprehensive approach than the corresponding manual workflows evaluated here.

## Acknowledgements

Support was provided by Hevolution Foundation to the Buck Institute for Research on Aging (HFPART-23-1422047) to LME, GL, JKA and JCN and NIH NIDDK 5R01DK135807 and a Sponsored Research Agreement from Calico Life Sciences LLC to AAG. An image of human pancreatic islets and some of the islets were provided by the NIDDK-funded Integrated Islet Distribution Program (IIDP) (RRID:SCR_014387) at City of Hope, NIH Grant # U24DK098085.

## Author contributions

Conceptualization: AAG, VMK. Human muscle study clinical concept: JCN, BJS. Respirometry and pancreatic β-cell study concept: AAG. Mitochondrial screening concept: AAG, VMK, JKA. Investigation and data analysis: VMK, SAW, CAL, VMD, EBS, ISB, HC, ST, JB, SG, SS, CS, ADC, SVA, EG, AAG. Funding acquisition: LME, GL, JKA. Supervision: JCN, BJS, SM, GL, LME, JKA, AAG. Project administration: LME, GL, AAG. Writing-original draft: AAG, VMK, SAW, VMD. Writing-review and editing: AAG, LME, JKA, VMK, SAW, VMD.

## Online Methods

### System Configuration

PickliPy was developed and validated on a Revvity Explorer G3 robotic workcell (Revvity, Waltham, MA) housed within a HEPA-filtered enclosure. The workcell integrated plate storage, liquid handling, and imaging components connected by a Plate::Handler^TM^ Flex (PFlex) robotic arm traveling on a linear rail. Plate storage comprised a LiCONiC STX44 robotic incubator (LiCONiC AG, Mauren, Liechtenstein) and a 63-position random-access rack. Labware handling included an Azenta 4AS plate sealer and XPeel seal peeler (Azenta, Burlington, MA) and an Agilent VSpin centrifuge (Agilent Technologies, Santa Clara, CA). Liquid handling was performed by three complementary systems: an Echo 650 acoustic dispenser (Beckman Coulter, Palatine, IL), a JANUS (Revvity) liquid handler equipped with single-tile Peltier heating/cooling devices CPAC Ultraflat Part No. 7000190 (Inheco GmbH, Martinsreid, Germany), and a MultiFlo FX (Agilent) bulk dispenser. High-content imaging was performed using a Revvity Operetta CLS spinning-disc confocal microscope. All instruments were coordinated by Revvity Plate::Works^TM^ scheduling software (version 6.30) interfacing with Echo Client Utility software (version 3.2.3). Supporting IT infrastructure included two image-analysis workstations with consumer-grade

GPUs, and a centralized file server. There was no laboratory information management system (LIMS) or other centralized high-throughput-screening-specific software or service used.

Chemical library data were handled with Microsoft Excel. Barcode readers in the workcell or in the Echo 650 were not used. Barcodes below refer to labels programmatically assigned to plates, tracking each plate from picklist design through merging metadata with analysis results. All shown experiments used the above workcell fully integrated solely through Plate::Works^TM^.

All dispense events within an experiment, spanning multiple source and destination plates were compiled into a single picklist by PickliPy, and the scheduling software automatically switched plates in the dispenser when needed.

### Materials

For the Echo 650 either Echo® qualified 384-well polypropylene 2.0 (384PP; Beckman Coulter) or cyclic olefin copolymer low dead volume (384LDV) microplates were used as source plates. For assay-ready storage plates 96-well V-bottom polypropylene plates (Corning, Corning, NY; #3363) were used. All other ADE experiments used 384-well glass-bottomed plates (Cellvis #P384-1.5H-N). The manual culturing of dispersed human pancreatic β-cells used glass-bottomed 96-well half area plates (Corning #4580). The small-molecule library was from TargetMol, Mitochondria-Targeted Compound Library 950 (#L5300); and it was extended with compounds of interests (including MIC (Coumarin 478; Cayman Chemical, Ann Arbor, MI; #40754) and UA (Sigma-Aldrich, St. Louis, MO; #SML1791) reported here). The TargetMol L5300 library entry “AMPK Activator 2 hydrochloride” was assigned as the parent structure 1-isobutyl-5-(4-trifluoromethylphenyl)biguanide (TargetMol T60675; CAS 2410961-69-0) by matching the supplier-provided structure to compound 7a in Xiao et al.^46^; chemical identity was not independently confirmed analytically. HEK293FT cells were from ATCC (Manassas, VA). The plasmid construct pcDNA3.1-mito-AT1.03 was a kind gift from Hiroyuki Noji, Osaka University. The Matrigel used was hESC-Qualified Matrix, LDEV-free (Corning #354277). The Geltrex was the Reduced-Growth Factor Basement-Membrane Matrix, LDEV-free, stem-cell qualified version (Thermo Fisher Scientific, Waltham, MA; #A1413302). Bovine serum albumin (BSA) was from Sigma-Aldrich (#126609). Trifluoromethoxy carbonylcyanide phenylhydrazone (FCCP) was from Cayman Chemical (#15218).

### Picklists and metadata

The picklist is a tabular data file following the specifications of the Echo 650 acoustic dispenser, listing individual well-to-well dispenses and their volumes for all microplates in the experiment, with plates identified by their barcodes. Inventory and process files are tabular data files defining microplate storage locations and the order of processing specifically for Plate::Works^TM^. Metadata generated by PickliPy documents the planned dispense events and is available for association with analysis data. Generated plate maps can be copied and pasted to the Harmony (Revvity) image acquisition control and analysis software. Well-compound associations or merge files can be merged into library data in DataWarrior^18^, or applied to image data sets in Image Analyst MKII (Image Analyst Software, Novato, CA).

### Development of PickliPy

PickliPy was originally developed in Mathematica 13.3 (Wolfram Research, Champaign, IL), and then translated to Python 3.12. PickliPy is a command-line tool, with most of its input loaded from a single design file that is in Microsoft Excel format. The design file contains source (SRC), destination (DST), and optionally library (LIB) and destination blacklist (DST_Blacklist) worksheets, and additional calculation worksheets may be present. The script logic follows features and constraints of the scheduling software and the acoustic dispenser. An important constraint is that both source and destination plates are sequentially used in a predetermined order (process files), and a plate that exits the pipeline cannot be used again. Plate::Works^TM^ filters the picklists for barcodes, therefore, all dispense events for all plates may be placed in a single picklist, and Plate::Works^TM^ automatically switches plates when all dispense instructions for the current plate barcode have been completed. The acoustic dispenser has limitations on minimum and maximum well volumes, therefore, precise tracking of original volumes and expected use was implemented. The Echo 650 instrument can survey (measure volumes of) source wells manually, or through a survey command from the scheduling software. Therefore, original volumes are user-provided on the SRC worksheet, and optionally, if an Echo survey file (*platesurvey.xml) is present, the user-entered values are automatically updated. It may happen that a source well has insufficient volume to complete the experiment. To avoid skipping these dispense events, PickliPy switches to a new well of the same stock if available, based on the matching compound name and stock concentration on the SRC worksheet (well pooling). An error is signaled if no backup well is available. The code ensures dispensing accuracy by rounding transfer volumes to 2.5 nL increments (specifically for the Echo 650-series) and flagging potential rounding errors at low-volume dispenses for user review. As a workaround for the limitation of a maximum 500 nL dispense volume from low-dead volume (LDV) source plates, larger dispenses are automatically split into up to 500 nL blocks. The script optimizes dispensing order for the shortest path for plate movements.

The DST worksheet is structured to provide values for variables (e.g. assay well volume), provide plate and concentration maps, and label definitions. The user enters arbitrary labels to the plate map, and a label definitions list below the map to associate these labels with compound names and final concentrations. One label can be associated with multiple compound names for combinatorial dispense, in consecutive rows of the label list. Furthermore, each plate map cell may contain multiple labels. The SRC worksheet associates compound names with source plate positions, stock concentrations and available volumes. Plate maps and the following label lists may be repeated in the same worksheet. If the same picklist name is reused, these will be merged in the same picklist.

### Pre-dispense validation

Before generating a picklist, PickliPy verifies that all compound labels in the destination layout are defined in the source inventory, that no source well definitions are duplicated, and sufficient volume is available to perform all dispense events. Dead volumes are specified by the user in the SRC worksheet and must accurately reflect the source plate type to ensure correct volume predictions.

### Randomization

PickliPy.Assay optionally randomizes the dispense wells, maintaining layout boundaries and sections, and attempting to maximize the distance between replicate wells and avoid adjacency. All generated metadata reflect the final, randomized dispense locations.

### The assay picklist generator

PickliPy.Assay supports well-by-well control of dispense based on one or more plate maps of labels and concentrations. The DST worksheet plate maps allow users to specify compounds and their concentrations in the following ways: 1) one or more labels per worksheet cell (where labels define compound stocks and final concentrations); 2) one or more compounds with concentration per label definition, or 3) a single label per cell plus a concentration map to allow setting numeric values for concentrations and using Excel autofill or formulas. In the DST worksheet, each plate map describes dispensing from one specific source plate to one or more destination plates. Merging multiple plate maps allows 2-dimensional titrations, and titrations in the presence of other added compounds, and dispensing from multiple source plates into a single destination plate.

### The screening picklist generator

PickliPy.Screen supports library screening with library reformatting, control addition, multi-dose dispense, and blacklisting faulty assay wells. The script threads a standard library table (LIB worksheet) into a plate map template, using as many destination plates as needed to dispense the whole library. The user creates a single destination plate template (DST worksheet) that defines where library compounds and controls will be dispensed. Controls are defined similarly to PickliPy.Assay with a label associated with a compound and its concentration, and these control compound names are listed in the SRC worksheet. In contrast, library compounds are not named, but defined by label placeholders (e.g., #1, #2, #3) to mark layout positions. Label placeholders in turn are associated with compound name placeholders and their concentrations that are filled sequentially from the compound library (LIB worksheet) by the script. In the source worksheet (SRC), controls are explicitly listed, optionally with a wildcard (*) plate barcode, indicating that these compounds occupy the same well on every source library plate.

Library compounds are defined using a single template row with column references that pull compound names, wells, and plate barcodes from the LIB worksheet. This template row defines all compound locations in the screen other than the defined controls above. In the DST worksheet, well replicates are created by repeating a label in the layout, while multiple doses are created by using different labels with the same compound name placeholder at different concentrations given in the label definitions below the plate map layout. Here, the final concentrations may be referenced to a LIB worksheet column, so each compound in the library can be used at its own predetermined concentration.

### The picklist generator for mitochondrial membrane potential assays

PickliPy.BlueTable is a specialized version of the assay method for making assay-ready plates for the mitochondrial membrane potential assay we previously described^32^. This method calculates dispense volumes and matches the well pattern defined in the potentiometric assay dilutions table^32^. This table lists 3 consecutive additions for time courses in a 3×6 pattern in partial 96-well format with desired treatments for each well as final and stock concentrations, including combinations of treatments. The compound stocks are dispensed into assay-ready plates for making 2×-concentrated additions in assay buffer using the carry-over dilution paradigm that enables keeping the concentration of previously added treatments constant during consecutive half-replacements of assay buffer^32^. The dispense is from a “tool library” (source) Echo-qualified microplate that is hand-filled with the required compounds, matching the inventory. This “tool library” plate also serves as storage for the compound stocks. The modifications to PickliPy.Assay dilutions table was to separate the inventory table (SRC worksheet) into a different file and allow for definitions of combinations and vehicles. Each design file describes an individual customized assay-ready plate and PickliPy.BlueTable converts a batch of such design files into a single picklist. The volume usage of the source plate is tracked between subsequent batches by the survey function of the Echo 650 instrument. On the day of the live-cell experiment, the user thaws, spins, and unseals the assay-ready plate, dissolves the dispensed reagents in assay medium, and transfers them into the assay reservoir.

### Post-dispense quality control

The Echo 650 generates log files for each operation: print files record transfer events including target volumes, actual volumes dispensed, and error codes for skipped wells while survey files record measured well volumes. For screens and experiments spanning many plates, manually reviewing these logs is impractical. PickliPy.QC provides a graphical interface that scans the Echo log directory for recent XML files, displays extracted plate barcodes for identification, and allows batch processing of selected files. For each transfer, the tool reports source well, destination well, compound identity (matched from the library database by plate barcode and well position), target volume, actual volume dispensed, and any reason for a no-dispense error. Volume discrepancies and dead volume violations are calculated internally and flagged only when present. Output is consolidated into a single Excel workbook. If any wells were missed or incompletely dispensed, PickliPy.QC may be used for generating a corrective picklist containing only the failed transfers, which can be loaded directly into the scheduling software for a targeted re-run without error-prone manual picklist editing.

### Agentic use of PickliPy

Agentic use of PickliPy, supporting creating or troubleshooting design files for PickliPy.Assay and Screen were tested in Visual Studio Code with OpenAI Codex extension. To this end the PickliPy git (see below) was cloned and opened as a folder in Visual Studio Code. Then, Codex was prompted to use “picklipy-excel-design-skill” to make or troubleshoot design files using PickliPy.Assay or Screen. See example prompts in Fig. S8.

### Plate::Works^TM^ automation with PickliPy and downstream use of metadata

In Plate::Works^TM^ a simple one-thread assay with two plate types (source and destination; Fig. S4c, left) or a two-thread design, one thread for each plate type (Fig. S4c, right) were used for dispensing into empty storage plates, making assay ready plates or into live cell cultures.

PickliPy-generated picklist files were used in the ECHO Pick function “Picking list” parameter, while the inventory and process files were used in the “Method/Entry” tab configuring the storage in the INCUBATOR and in the RACK components with the READ_INVENTORY function, “Allocation list” and “Process list” parameters. Plates were positioned manually in the LiCONiC incubator or in the random-access rack to match the inventory files that always followed alphabetic ordering of barcodes assigned in the PickliPy design file. The merge table file was used in Image Analyst MKII, after opening the data set exported from harmony with the “Write Archive” option. To this end, in the Multi-dimensional Open dialog, Position Names tab, “Load position names from well-plate-compound list” command was used. In Revvity Harmony, PickliPy-generated plate maps can be entered in the Assay Layout Editor using copy and paste.

### Dispensing on blotting paper with ADE

To dispense liquid with ADE on paper we designed a microplate-format blotting paper adaptor, and single-material extrusion 3D-printed it using PETG (Polyethylene Terephthalate Glycol) as filament material with a MakerGear M3-id 3D-printer (MakerGear, Beachwood, OH). Meshes were deposited at https://3d.nih.gov/entries/3DPX-023601.

### Comparison of aqueous and DMSO ADE protocols

Comparison of acoustic dispensing of dilute and concentrated, aqueous and DMSO solutions was performed for each solvent using both 384PP_DMSO2 and 384PP_AQ_SP2 dispense protocols of the Echo 650 instrument. The dispense was executed into 384-well glass bottom plates (Cellvis #P384-1.5H-N) containing 50 µL H_2_O, and fluorescence was quantified using a Pherastar FS (BMG Labtech, Cary, NC) microplate reader (not integrated in the workcell).

Fluorescein (240 µM final) was added to H₂O (100%), DMSO (100%), and aqueous solutions of glucose (3 M), succinate (1 M). Each dispense was performed with 2 plate replicates, including the manual pipetting, and the dispenses were run in triplicate. As a control, the stained solutions were also manually diluted in deep-well 96-well plates and subsequently transferred to the same type of destination plate to replicate the exact format used with ADE. To compare slopes, an unweighted linear-regression fit was performed with fluorescence as the response and dispense method, nominal volume, and their interaction as predictors. Differences in calibration behavior were assessed by comparing slope ratios relative to manual dispense using 95% confidence intervals.

### Evaporation and moisture absorption during ADE

Evaporation tests were performed by manually pipetting 50 µL of H_2_O into each well of a 384 well PP plate. Moisture absorption was performed the same way as the evaporation experiment but using 50 µL fresh anhydrous DMSO. This plate was delivered into the source position on the Echo 650 and surveyed using the Echo Client Utility software. The first plate survey is the starting baseline read, time 0 min (T0). Thereafter, the plate remained in the Echo 650 and was surveyed at 30 min intervals up to 120 min (T120). These data were used to determine the rate of evaporation over time. To ascertain whether the rate of evaporation and the position of the door affect changes in volume, this experiment was conducted with both the instrument’s door open and closed.

In longer experiments with aqueous solutions, substantial evaporation can occur due to the air flow through the dispense chamber of the Echo 650; therefore, the minimum source well volume indicated in the SRC worksheet may be set higher than the instrument’s cutoff value.

Separating repeated dispense events (e.g. during a long time course) into multiple picklist files allows scheduling software to eject and lid the source plate during incubation periods (Fig. S4c).

### Isolation of human skeletal mitochondria

Muscle biopsies were collected from subjects recruited by the Human Biology Core at the Buck Institute by a licensed physician trained in the procedure under sterile conditions. Written informed consent was provided ahead of any study procedures, and the study was conducted in accordance with the current Declaration of Helsinki (approved by Advarra IRB: Pro00078112, BUCK_2401, An observational study of aging biomarkers in older elite athletes compared to non-exercising healthy older adults, registration number: NCT06540287). Samples and associated data were coded by the clinical team at the point of collection. Investigators performing the molecular/analytical work had access only to coded samples and were blinded to the identity key, which was retained solely by the clinical team. Participants followed a standardized pre-biopsy protocol (2 h fast except water/caffeine, 12 h alcohol/cannabis abstention, 24 h exercise restriction, 7-day NSAID/aspirin washout, and a standardized oatmeal meal ≥2 h prior) to minimize acute metabolic confounds on the collected tissue.

The subject data presented were a subset used for benchmarking in an ongoing clinical study comparing athletes and unexercised individuals. The total subject count was n = 38, sex distribution was 50% female, mean ± SD age was 71.5 ± 4.0 y, mean ± SD BMI was 23.0 ± 3.4, activity distribution was 89% athlete. Of these subjects, only those 30 were included into the coefficient of variation analysis (Fig. 3d-g) where all n = 4 replicates (2 replicate wells on each of 2 experimental plates) passed instrument QC based on baseline stability and trace quality.

Compound dosing was performed based on availability of surplus sample, therefore replicate numbers varied.

Biopsies were obtained from the right vastus lateralis under local anesthesia (1% lidocaine, ≤4 mg/kg, up to 200 mg, without epinephrine) using a 5–6 mm Bergstrom Needle under suction, yielding 3–5 consecutive tissue sections per procedure (≥200 mg total; a second pass was performed only if this threshold was not met).

Tissue was placed in 1× phosphate buffered saline (PBS) on ice. A 3×3×5 mm piece was cut into partitions along the fiber grain to preserve fiber length. A 30-50 mg (wet weight) sample was placed into 500 µL Chappell-Perry buffer 1 (CP1) (50 mM Tris, 2 mM EGTA, 100 mM KCl pH-adjusted to 7.4 by HCl) (all from Sigma-Aldrich) and transferred to ice for mitochondrial isolation.

Mitochondrial isolation and assaying were based on previous work^19^ with modifications. All manipulations were performed on ice or at 4 °C. Surgical scissors were used to roughly chop biopsies in 500 µL Chappell-Perry (CP1) buffer. The biopsies rested for 2 min before carefully removing the CP1. An amount of 300 µL of CP2 buffer (CP1 supplemented with 5 mM MgCl2, 0.5% w/v BSA, 1 mM ATP and 2.5 U/mL of Protease Type VIII (Sigma-Aldrich, #P5380)) was added along with a stir bar. The sample was pressurized in an ice-filled chamber up to 800 psi in N_2_ gas over a magnetic stirrer over the course of 1 min and cavitated for 10 min before rapid decompression. Immediately 1 mL of CP1 was added. The sample was transferred using pipette tips cooled in 0.5% BSA-containing CP1 to a handheld homogenizer. The sample underwent 10 strokes (2 mL loose pestle B) before the homogenate was transferred to a clean 2 mL Eppendorf tube. The homogenizer was rinsed with 200 µL CP1 and transferred to the collection tube. The tube was centrifuged (4 °C, 500 g, 10 min, accel:3, decel:3). Supernatant was collected and filtered through a 40 µm filter before being centrifuged (4 °C, 10,000 g, 10 min, accel:3, decel:3). Supernatant was again discarded, the pellet was gently washed with CP1 and resuspended with 40 strokes in 1 mL CP1, diluting to 2 mL after resuspension. The sample was centrifuged (4 °C, 10,000 g, 10 min, accel:3, decel:3), supernatant was discarded, pellet was gently washed in CP1 and resuspended in 100 µL of CP1. Yield was normalized to the result of Pierce BCA protein assay (Thermo Fisher Scientific, #23225), typically measured between 60-200 µg.

### Dispense of Seahorse XF kit plates for deep bioenergetic phenotyping

The liquid handling burden associated with filling the Seahorse XFe96 assay wells and cartridge addition ports was mitigated by ADE and manual pre-dispensing assay-ready “kit” plates. This was achieved by separating ADE-compatible, small volume additions (DMSO or aqueous additions up to 7 µL, typically 100 nL) and bulk aqueous volume additions (assay buffer + substrates, nucleotides, etc.) into four plates (AQ Port, AQ Substrate, ECHO Port and ECHO Substrate; see below), each stored frozen until use. The “substrate kit” is added to the assay wells as the basal medium, while the “port kit” is added to the addition ports of the Seahorse XFe96 cartridge after mixing the ADE and bulk dispensed volumes (Fig. 3b). Dispensing relied on PickliPy.Assay, using additional worksheets in the design Excel workbook to perform the required calculations (e.g. standard Seahorse XF port addition dilution calculations^19^)

### Seahorse XF respirometry on isolated mitochondria

A Seahorse XFe96/XF Pro sensor cartridge (Agilent) was hydrated the night before using Seahorse XF calibrant at 37 °C. Seahorse XFe96/XF Pro cell culture microplates were degassed at 37 °C in ambient air for at least 4 h. Port and substrate assay-ready plates—each consisting of 1 aqueous (AQ) plate and one acoustically dispensed (ECHO) plate, 4 plates total per sensor cartridge were thawed and spun (4 °C, 500 g, 3 min, accel:1, decel:1). 250 µL of the AQ Port plate was transferred by rows using a 12-channel pipette to the ECHO Port plate (rows A, B, C, D into rows E, F, G, H respectively) dissolving its contents. 20 µL of this mixture was dispensed by rows using a 12-channel pipette into the ports of the sensor cartridge (rows E, F, G, H of the kit plate into ports A, B, C, D respectively, for every row of the assay plate), filling all 384 ports.

Thus, each well in a column of the 96-well assay plate received the same four port additions. The filled plate was calibrated in the Seahorse XFe96 Analyzer. Meanwhile, mitochondrial suspension was plated into the degassed and cooled XFe96/XF Pro cell culture microplate (without any coating) at 1 µg/well in 20 µL of MAS1 buffer (2 mM HEPES, 10 mM K_2_PO_4_, 1 mM EGTA, 70 mM sucrose, 220 mM mannitol, 5 mM MgCl_2_, pH-adjusted to 7.4 at 37 °C using KOH) (all from Sigma-Aldrich) supplemented with 0.5% w/v BSA, and the same buffer was added into background wells, and the plate was centrifuged to immobilize mitochondria (4 °C, 2000 g, 20 min, accel:1, decel:3).

250 µL of the AQ Substrate plate was transferred by rows using a 12-channel pipette to the ECHO Substrate plate (rows A, B, C, D into rows E, F, G, H respectively). 100 µL of room temperature substrate was gently added using a 12-channel pipette to the cell culture/assay plate over the mitochondria (rows E, F, G, H into rows A/E, B/F, C/G, D/H respectively). The substrate conditions varied well-by-well with technical duplicates, and their combinations with the above column-wise differing port additions created 22 different duplicate assay conditions and four background wells (where no mitochondria were added) in each half of the assay plate.

For compound dosing experiments (UA, MIC, Azure B), 120 µL of room temperature substrate was mixed with an ADE drug plate which contained 4 rows of drug. 100 µL of this was then added to rows E-H of the cell culture plate, in the rows below the non-drugged substrate added to rows A-D of the cell culture plate.

After addition of substrates, the assay plate with the mitochondria was immediately loaded into the XFe96 instrument. Settings for the assay run were the same for each of the 5 measurement segments (baseline, port A, port B, port C, port D), consisting of 3 cycles of 1 min of mixing, 1 min of waiting, and 3 min of measuring.

### Measurement of the P/O ratio

The P/O ratio (or ADP/O ratio; the number of ATP molecules synthesized per consumed oxygen atom) was calculated from ECAR and OCR, based on previous work from our group^20,22^ and others^21^. Buffering power was calculated as the slope of pH over HCl (mM) from a dedicated duplicate experiment in the Seahorse XFe96 Analyzer. To this end, 4 additions of 20 µL of 2.4 mM HCl were made into 120 µL MAS1. Experimental ECAR was converted to proton production rate (PPR) by dividing by the measured buffering power (0.32 pH/mM) and multiplying by 2.28 µL (the microchamber volume) and a scaling factor of 1.6 (provided by Agilent) in Fig. 3g (Raw P/O). In subsequent data (Fig. 3h), raw P/O was further corrected by multiplying 1.31 to match the theoretical succinate and glutamate + malate values. The P/O ratio was calculated as - PPR/(2*OCR) using oligomycin-sensitive rates, thus its value was independent of proton leak. Oxidation of succinate or glutamate + malate does not result in net proton production or capture other than by phosphorylation, avoiding a distortion of the alkalinization-derived P/O.

In contrast, other commonly used substrate combinations, such as pyruvate + malate or palmitoyl-carnitine + malate may result in carbonic acid or citrate formation, depending on the completeness of oxidation in the TCA cycle. To track this, we calculated alkalinization derived P/O values in Table S1.

### Calculation of theoretical values of alkalinization-derived P/O ratio

Regular P/O values shown in Table S1 were calculated by standard chemiosmotic stoichiometry: electron entry determines pumped protons, pumped protons are converted to ATP equivalents, and substrate-level phosphorylation and carrier costs are added or subtracted. To this end, a stoichiometry of *H*^+^/*ATP* = 11/3 was used, based on an 8-c-subunit mammalian ATP synthase ring plus 3 carrier protons per 3 ATP^20^. For alkalinization-derived P/O the ATP-equivalents were corrected for acid-base proton production. To this end scalar H^+^ yield (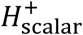; as given by the atom- and charge-balanced net ionic equation in Table S1), and CO_2_ and citrate formation were considered. Respiratory CO_2_ was treated as contributing to measurable acidification only to the extent that it equilibrates to bicarbonate: 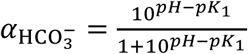 with *pH* = 7.4 and *pK*_1_ = 6.093, giving 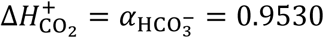. Thus, each CO_2_was counted as releasing 0.953 H⁺. Citrate was handled separately because exported citrate at pH 7.4 is not exactly citrate^3−^. The average number of deprotonated carboxyl groups per citrate molecule at pH 7.4, treating citrate as a triprotic acid using pK_a1_=3.13, pK_a2_=4.76, pK_a3_=6.40 and 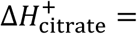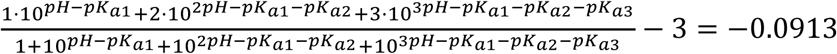, meaning that pH-equilibrated citrate consumes about 0.091 H⁺ per citrate relative to the formal citrate^3−^. The alkalinization-derived ATP-equivalent yield was then: 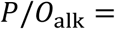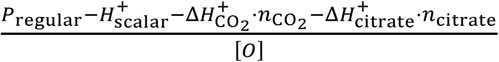.

### Human primary dispersed pancreatic islet cell cultures

Dispersed islet cell cultures were prepared from post-mortem human islets from multiple islet centers; Prodo Laboratories (Irvine, CA), University of California San Francisco Diabetes Center and through the Integrated Islet Distribution Program (IIDP). All samples were deidentified. Type 2 diabetic (T2D) organ donors were defined by the islet center based on HbA1c and medical history. Islet cells were dispersed using AccuMAX, cultured in Prodo PIM(S) medium and plated in PEI (1:15,000 w/v; overnight, 37 °C)-coated glass-bottomed half-area 96-well microplates (Corning #4580) at 5,000 cells/well in the presence of 5% v/v Geltrex as previously described^30^ for 6-10 days. Each plate received cells in 18 wells in a 3×6 pattern. The growth medium for dispersed islet cultures was PIM(S) supplemented with human AB serum (5%, Corning #CV35-060CI or Gemini #100-512 or #100-712), glutamine plus glutathione (PIM(G); Prodo) according to the supplier’s recommendations, plus 100 units/mL penicillin and 100 µg/mL streptomycin.

Data from total 32 islet donors were used in this study, recorded between 2017 and 2026. FCCP data are shown for 12 islet donors assayed after converting our liquid handling workflows to ADE (age = 50.9 ± 2.1 *y*, BMI = 27.0 ± 0.9 , HbA_1c_ = 5.6 ± 0.27 ; mean ± SE; 83 % male sex) and for the last 12 islet donors assayed using manual pipetting before this (age = 49.5 ± 2.7 *y*, BMI = 28. ±1.4, HbA_1c_ = 5.73 ± 0.16; mean ± SE; 75 % male sex). For Atpenin A5 dose-response, all islet cell preparations assayed in this time range for the corresponding conditions were included (n = 20 subjects). These subjects were split in an older (age = 56.8 ± 1.4 *y*, BMI = 28.7 ± 1.3 , HbA_1c_ = 6.33 ± 0.35 ; mean ± SE, n = 10; 90 % male sex) and a younger (age = 40.6 ± 1.7 *y*, BMI = 28.1 ± 1.9 , HbA_1c_ = 5.62 ± 0.16 ; mean ± SE, n = 10; 40 % male sex) group. Detailed data are in Table S3.

### Plasma and mitochondrial membrane potential assay on pancreatic β-cells

Absolute mitochondrial membrane potential (ΔψM) was measured by fluorescence microscopy using tetramethylrhodamine methyl ester (TMRM) and FLIPR (#R8126 FLIPR Membrane Potential Assay Explorer Kit; red version; Molecular Devices) as previously described^30,32^ and with modifications to the assay buffer preparation to use ADE. Recordings were performed on two Nikon Eclipse Ti Perfect Focus System fully motorized wide-field fluorescence microscopes that were equipped with custom Lambda 821 LED light sources (Sutter Instruments, Novato, CA) or earlier with a Lambda LS Xe-arc light source, with Lambda 10-3 emission filter wheels, an Andor iXon Life 888 (Oxford Instruments, UK), or a Cascade 512B (Photometrics) EMCCD camera and a Nikon or an MS-2000 linear encoded motorized stage (ASI; Eugene, OR) with NIS Elements 5.20 (Nikon, Melville, NY) using an S-Fluor 10× air lens. TMRM and FLIPR signals were collected at 100 s intervals, using the following filter sets, given as LED nm, excitation –– emission in nm/bandwidth, for TMRM: 561, 586/20 – 641/75 (30 ms exposure time, 14% power) and for FLIPR: 506, 509/22 – 542/27, using a 459/526/596 beamsplitter (30 ms, 7%; all from Semrock, Rochester, NY). Cells were washed three times 4 h before recording with a modified culture medium (non-fluorescent RPMI with 3 mM glucose, 1 mM NaHCO_3_, 20 mM TES, 2 mM glutamine) supplemented with 7.5 nM TMRM, 1:100 FLIPR, 1 µM Na-tetraphenylborate, 1 µM zosuquidar. The cells were incubated in an air incubator at 37 °C to equilibrate the probe and were washed another two times 30 min before the baseline recording.

For stimulations and challenges, assay-ready plates were pre-made using PickliPy.BlueTable and stored at -20 °C. To this end, 100 nL DMSO stock was dispensed per well of a V-bottom 96-well polypropylene plate by ADE. Dispensed volumes were made up by one or more compound stock plus DMSO, allowing dose-responses and combinatorics to be performed at constant vehicle concentration. About an hour before the recording, the contents of each well of the V-bottom plate were dissolved in 200 µL assay medium to yield 2×-concentrated treatment. Dissolved stocks were immediately transferred into 700 µL glass inserts (to prevent depletion of TMRM by the plastic sidewalls) in a deep-well 96-well reservoir to be manually pipetted during the recording. In contrast, the earlier manual version of this assay hand-pipetted 0.35 µL DMSO stocks into the glass inserts before adding the 700 µL assay medium, and dose-response recordings used appropriately diluted stock solutions for each final concentration. Using PickliPy.BlueTable and the original stock dilution design tables an important feature of the original paradigm was preserved that the successive additions do not dilute previously added modulators, due to a carry-over step during making the reservoir plate^32^. The time-course recordings comprised of baseline, treatment by manually half-replacing medium using a multichannel pipette, recording, second treatment and recording and additions for the potentiometric calibration as described earlier^30,32^. Recordings were analyzed in Image Analyst MKII. ΔψP and ΔψM were calibrated using the “Complete Calibration with known kP and Goldman potentials using Neural Network” and “Complete Calibration” paradigms. Beta-cells were identified by dithizone staining and shown data represent only β-cells and not all dispersed islet cells.

### Analysis of ΔψM in response to bioenergetic challenges

Millivolts-calibrated membrane potential data were exported as GraphPad Prism files from Image Analyst MKII, and imported into Mathematica 13-14 (Wolfram Research, Champaign, IL), using a custom XML parsing code. Here a database was created holding all compound addition-evoked potential changes using the Dataset function. To this end metadata describing additions were originated from the compound names and concentrations set up in the BlueTable design files and propagated through Image Analyst MKII and Prism export to Mathematica, and metadata describing donor clinical values and experimental parameters were added. This database was then used for descriptive statistics, for multivariate regression using the ResourceFunction["BayesianLinearRegression"], and for dose-response analysis. To analyze ΔψM depolarization in response to increasing SDH inhibition with Atpenin A5, a constrained, L1-regularized nonlinear Hill model was used. For subject *i*, the response at inhibitor concentration *c* was modeled as 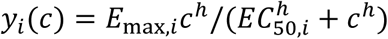, where *E*_max,*i*_ = exp(*μ_E_* + *d_E_*_,*i*_) and *EC*_50,*i*_ = exp(*μ_C_* + *d_C_*_,*i*_) and *μ* terms are cohort-level intercepts and *d* terms are subject-specific deviations on the natural-log scale. A single Hill coefficient, ℎ, was shared across the cohort to reduce model dimensionality and limit the strong covariance among *E*_max_, *EC*_50_, and slope that is expected when each subject is represented by only four concentrations and the maximal response may not have been reached. The lower asymptote was fixed at zero, because responses were expressed relative to matched vehicle controls, and occasional negative responses were treated as measurement variation around the null response rather than as evidence of a nonzero subject-specific baseline. All observations were fitted simultaneously by constrained nonlinear least squares, with ℎ > 0 and subject-specific *EC*_50_ < 0.5 µM (maximal used Atpenin A5 concentration).

### Generation of a HEK293 line stably expressing the mito-AT1.03 ATP-sensor

The ORF of mito-AT1.03 was PCR amplified from the original pcDNA3.1 plasmid^50^ and cloned into pENTR-D-TOPO following the manufacturer’s instructions. Using LR-recombination with a pLenti-CMV-Hygro plasmid, pLenti-mito-AT1.03 was generated and verified by sequencing.

Lentivirus was generated with the ViraPower (Thermo Fisher Scientific) third-generation lentiviral packaging kit in HEK293FT cells and concentrated by ultracentrifugation. To generate a mito-ATeam-expressing cell line, fresh HEK293FT cells were transduced at a low MOI and cultured under hygromycin selection. Then, cultures were sorted for medium-high expressors using a BD FACSAria II cell sorter (BD Biosciences, San Jose, CA). HEK-mAT1.03 cells were cultured in DMEM (high glucose with glutamine) supplemented with 10% FBS plus 100 units/mL penicillin and 100 µg/mL streptomycin.

### Matrigel coating microplates for HEK-mAT1.03 screening

Cellvis P384-1.5H-N 384-well glass-bottomed microplates were degassed for 1 h in the Revvity workcell’s LiCONiC incubator to minimize the likelihood of bubble formation during the Matrigel gelling phase. Matrigel was thawed on ice and diluted according to the official recommendation using the same DMEM culture medium used for mito-ATeam cell culturing. Before starting the automated Matrigel coating protocol, tips for liquid dispensing were pre-cooled at 4 °C for at least 30 min and 3 Peltier cooling tiles were positioned on the JANUS liquid handler’s deck and set to cool to 0 °C for at least 20 min. Peltier cooling tiles were used to keep Matrigel and PBS cool in large 1-well reservoirs, and to cool the microplates. During the coating protocol, microplates were brought one at a time from the incubator to cool on a cooling tile of the liquid handler’s deck for at least 3 min, while tips were kept cool submerged in Peltier-chilled PBS. Using the liquid handler’s 96-channel MDT arm, plates were coated at 12 µL per well, for the inner 240 wells of the plate. Immediately after the liquid dispensing process, plates were centrifuged using the VSpin at 500 g for 90 seconds at room temperature, left for 3 min on a rack at room temperature to slowly warm up, and then brought back to the incubator at 37 °C to incubate overnight before use in cell seeding. If coated plates were not planned to be used within 24 h, they were only incubated for 30 min and manually stored in a 4 °C fridge until use. After incubation, medium was aspirated from the inner 240 wells before seeding cells.

### Small-molecule library handling

The Mitochondria-Targeted Compound Library was purchased in 384PP format and daughtered with direct transfer or with subsetting and reformatting for dose-responses into 384LDV plates using picklists generated by PickliPy.Screen for the Echo 650 instrument.

Within the Revvity G3 Explorer workcell, all plate movements were executed by the PFlex robotic arm under Plate::Works^TM^ scheduling software coordination. Thawed, sealed library plates were transferred from a random-access rack to a VSpin centrifuge to consolidate liquid at the well bottom. This is a requirement for accurate acoustic dispensing, as accurate meniscus detection and position are crucial for ADE. Library plates were then unsealed and transferred to the Echo 650 along with empty daughter LDV plates. For direct daughtering, the transfer was one-to-one: source plates containing 320 filled inner wells were replicated identically into daughter plates. For the primary screen, control compounds (undiluted DMSO, FCCP + oligomycin (4 mM + 2 mg/mL), myxothiazol + oligomycin (4 mM + 2 mg/mL), and CCCP (10 mM)) were added manually to each source plate to be distributed to daughter plates. For dose-response consolidation, the same automated Plate::Works^TM^ protocol was used, and source plates were sequentially and automatically exchanged in the Echo until all compounds of interest were transferred onto the single daughter plate based on the picklist created by PickliPy.Screen. Upon completion, all daughter plates were sealed and returned to the rack.

Dispense accuracy was validated post-run using PickliPy.QC (see “Post-dispense quality control”). PickliPy.QC generates an updated inventory sheet based on the daughter plate contents, which can be used directly as the LIB sheet input for subsequent screening picklist generation.

A standard 384-well library plate layout reserves columns 1-2 and 23-24 as empty (320 usable wells). For the mito-ATeam screen, this layout was reformatted. Each assay plate contained 75 compounds dispensed at three concentrations (1, 3, and 10 µM), along with 6 DMSO vehicle controls and 3 wells each of FCCP + oligomycin, myxothiazol + oligomycin, and CCCP as positive controls. Dispensing was restricted to the center 240 wells to avoid plate-edge interference with the water-immersion objective.

### Mitochondrial structure and [ATP] screen

HEK-mAT1.03 cells were cultured, drugged, and imaged in a custom non-fluorescent DMEM formulation (lacking phenol red, riboflavin, and folic acid) supplemented with 10% FBS (Thermo Fisher Scientific, #A5670701), 1% penicillin-streptomycin (#15070063), 20 mM TES, 1 mM sodium pyruvate, and 4 mM glutamine. Cells were seeded at 5,000 cells per well in 50 µL medium into the inner 240 wells (to avoid collision between the plate skirt and the lens) of Matrigel-coated glass-bottom 384-well microplates (see above). Each plate was seeded using the MultiFlo FX bulk dispenser with a 5 µL cassette peripump, then incubated overnight (37 °C, 5% CO₂) in the LiCONiC incubator to allow adherence. Compounds were then dispensed by the Echo 650 directly into the cell plates from the library daughter plates (384LDV) using picklists generated by PickliPy.Screen, with the entire library processed in a single batch. 50 nL was dispensed into 50 uL cell culture volume. These cell culture volumes in 384-well plates were safe in the inverted plate position during dispense. Then, the drugged cultures were incubated for 48 h. The seeding and compound treatment workflows were fully automated using the following devices: rack, incubator, lidder, centrifuge, seal peeler, plate sealer, MultiFlo FX dispenser, and the Echo instrument. These steps were controlled by Plate::Works^TM^ and the PFlex robotic arm.

Imaging was performed on the Operetta CLS using a 63× NA 1.15 water-immersion objective in confocal mode at camera binning 2. There were 16 fields acquired per well in a single focal plane using the Two Peak autofocus method, optimized for this cell line and plate type. Two channels were captured for ratiometric quantification of the mito-AT1.03 biosensor: CFP (excitation 435-460 nm, emission 470-515 nm, 80 ms exposure at 100% LED power) and CFP to YFP FRET (excitation 435-460 nm, emission 525-580 nm, 80 ms exposure at 100% LED power). The imaging workflow was fully automated using the following devices: incubator and microscope; controlled by Plate::Works^TM^ and the PFlex robotic arm.

### Mitochondrial morphomics

Image data were exported from Harmony (Revvity; “Write Archive” function) and the entire screen consisting of 13 plates (or 3 plates for the dose-response) was opened in Image Analyst MKII (Image Analyst Software, Novato, CA; “Open Instrument-specific TIFF folder” internal reader method). Compound identities and concentrations were mapped to well positions using the metadata file (table_merge.txt) generated by PickliPy.Screen, enabling automated annotation of all imaging data with treatment conditions. To measure all reported features, image-analysis pipelines (https://github.com/gerencserlab/IA-Mito-Morphomics) were executed in all wells of all plates as a “run in all positions” operation. The analysis was performed as an unsupervised, automated workflow without generating intermediate image datasets. Tiled frames were background and flat-field corrected and then stitched. Using the sum of the CFP and FRET images, mitochondrial profiles were generated by a combination of spatial filtering and morphological segmentation as previously described^31^ or using MoDL^43^. To adapt MoDL to our workflows, we improved MoDL’s patch handling to avoid edge artifacts, introduced resampling of images to match training and inference resolution and embedded MoDL in a Python Flask-based service for automation (https://github.com/gerencserlab/HTS-MoDL). Image Analyst MKII, probability maps from MoDL were post-processed by watershed segmentation attempting to discriminate between touching mitochondria. Cell boundaries were determined by Cellpose 3 segmentation using the cyto3 model in the ATeam sum images, were quality-controlled for sharpness based on punctate over diffuse index in the above sum image, and edge-touching cells were discarded. Cellpose was also used for automation by applying a Python Flask service (https://github.com/gerencserlab/IA-Cellpose-tools).

Morphological parameters were measured in whole view fields or in a hierarchical arrangement by converting QC-ed Cellpose cells to ROIs in Image Analyst MKII and first performing the given statistic (such as mean, variance or median of segment metrics or pixel intensity ratios) in each ROI and then a higher-level statistic on all ROIs of the view field. The following morphological metrics were calculated: 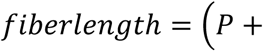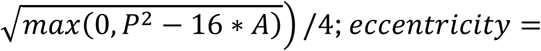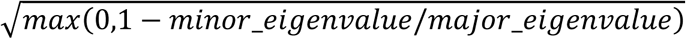, where P is the number of perimeter pixels and A is the area in pixels; minor and major eigenvalues are of PCA-based elliptic fits corresponding to the minor and major axes of the best-fit ellipse over the pixels of a segment. The skeletal length was the number of pixels per segment (mitochondrion) in a skeletonized image. Mitochondrial thickness evenness was quantified for each mitochondrial segment as the normalized Shannon effective number of skeleton-to-boundary distances. For distance values *x_i_* measured at *n* skeleton pixels, shares were defined as *p_i_* = *x_i_*/ ∑*_j_ x_j_*, Shannon entropy as *H* = − ∑*_i_ p_i_* ln *p_i_*, and thickness evenness as *E* = exp(*H*) /*n*. Values approaching 1 indicate uniform skeleton-to-boundary distances, whereas lower values indicate that the total distance is distributed less evenly along the mitochondrial skeleton. Mitochondrial skeletal length was expressed as ln(pixel area of the skeleton), because of the slightly better assay window as compared to not using logarithm. Notably, skeletal length resulted in discrete values of few pixels and this was not suitable for median and MAD (median absolute deviation) calculation. In contrast fiber length and to a greater extent eccentricity produced more granular data where both mean and median calculation were applicable.

ATeam FRET Ratio data were calculated per (visible unit of) mitochondria based on the mitochondrial profiles, by calculating first average fluorescence intensity over the profile and then calculating the FRET/CFP ratio. Then the mean, median or variance, of mitochondrial objects were calculated per view field, per cell or subcellular, using the above ROI definition. In some cases, means were Winsorized by capping values lower than the 1^st^ percentile or higher than the 99^th^ percentile. In addition, extreme-value distribution was described by calculating bottom or top (not shown) 5% share. Bottom 5% share was expressed as the fraction of the total signal that is carried by the lowest 5 percent of values, and was calculated as the sum of the smallest 5%-of-observations divided by sum of all values.

Morphology and ATeam data were recorded in the internal database of Image Analyst MKII and exported as CSV files for analysis in Python or in R. For benchmarking, the first 10 plates were used that were plated at identical cell densities.

### Assay Window Statistics

MAD-based Z′-factor was calculated as 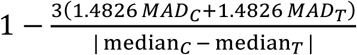, where C was for vehicle negative control (DMSO) and T for positive control and *MAD*(*x*) = median*_i_*|*x_i_* − median(*x*)|. The 1.4826 multiplier scales MAD to be equivalent to SD. To calculate dispersion of Z’-factors, a 20,000-resample well bootstrap approach was used calculating 95% confidence intervals.

### Training of MoDL mitochondrial segmentation model

Large view fields were first segmented with 4× upscaling using the original, super-resolution-microscopy-trained network of MoDL. The same view fields were also segmented by the above classical morphological segmentation algorithm. Objects detected by the morphological segmentation algorithm but not by MoDL were added to a new mask, forming the ground truth (or curated reference masks) for the training. The resulting mask and the original FRET+CFP sum images were randomly cropped to 512×512 patches. The 10 percent of lowest pixel entropy patches (of the ATeam image) and the 10 percent of lowest mask pixel count patches were discarded. This was performed on 2 DMSO, 2 CCCP, 2 myxothiazol + oligomycin - treated and 2 untreated wells, resulting in a total of 445 patches. Training was performed using default settings of the original repository in 50 epochs. The retrained network was then used without rescaling on our recordings.

All training and inference were performed on consumer-grade PC workstations or laptops running Microsoft Windows 11, using GPU computing with NVIDIA GeForce RTX 3090, 4090 or 5080 graphics cards.

### Hierarchical variance decomposition

Hierarchical variance decomposition was performed at the individual view-field level preserving the field-to-field replication structure before treatment-level aggregation for fiber length and ATeam FRET ratio variances. The observed total variance was measured in the population of mitochondria in the whole view field, the intracellular component was the median of ROI values measuring variance of enclosed mitochondria, and the raw cell-to-cell component was the variance of ROIs measuring median of enclosed mitochondria. For each field, the cell-to-cell component was corrected by subtracting the expected intracellular contribution per mitochondrion-per-cell that was calculated by dividing the raw value by the average number of mitochondria per cell. To test for significant effects, a two-stage cluster bootstrap (20,000 iterations) first sampled plate barcodes with replacement and then sampled wells with replacement within each selected plate. The same sampled wells were used for both variance components, preserving their pairing. Plates were considered as independent replicates and significance levels were calculated as BH-corrected FDR q-values.

### Pre-screen quality control with blacklisting

As part of the above workflow, after 24 h culturing, all microplates were imaged at 10× magnification using digital phase contrast on the Operetta CLS. Cells were counted using the Harmony software, and wells with plating defects were excluded from the screen using the blacklist feature of PickliPy.Screen.

### Dose-response follow-up

Selected compounds from the primary screen were tested in nine -point dose response (0.09–10 µM or 0.9–100 µM). A new subset daughter plate was prepared from the original 10 mM or 100 mM stocks alongside 33-fold intermediate dilution wells. The dual-concentration configuration exploited the large dynamic range of ADE to span the full dose range. Dose-response compounds were reformatted using picklists generated by PickliPy.Screen (see above, “Small-molecule library handling”). Data analysis was performed in Python on tabular output from Image Analyst MKII. All features were normalized to per-plate DMSO median fold-change to account for plate-to-plate variability in baseline signal.

### Dose-resolved co-response hierarchy analysis

Single-cell ATeam FRET ratio and mitochondrial morphology features were generated in Image Analyst MKII using the above-described “Mitochondrial morphomics” workflow, with per-cell rather than well-level output exported for downstream analysis. For the dose-response co-response hierarchy analysis, per-cell features were aggregated to well-level summaries using P1-P99 Winsorized means (P, percentile) and standardized against plate-matched DMSO vehicle controls. Within each compound and candidate concentration regime, local dose-response slopes were estimated separately for the ATeam FRET ratio (A) and each of the three morphological features (M) using ordinary least-squares models of each endpoint as a function of log10 concentration, with plate included as a blocking factor.

To screen for range-dependent or nonlinear behavior, full-range and auto-optimized breakpoint-defined low/high concentration regime were compared as exploratory models. The breakpoint was defined based on Bayesian information criterion improvement. Evidence levels were assigned by a gate hierarchy incorporating both-endpoint dose response (in-family FDR q-values < 0.05), cell-count-based viability (> 25%), denominator strength (for ATeam;|t| >= 3.2), one-slope adequacy by lack-of-fit testing (p ≥ 0.05), plate consistency (whether the effect reproduces across plates, requiring slope-sign agreement in at least two of three plates), saturation/floor-ceiling QC (always satisfied, not shown), and bounded Fieller confidence intervals for the local slope ratio. The A-M proportionality estimate was expressed as the local slope of the morphology metric (as a function of concentration) divided by the local ATeam FRET-ratio slope, with uncertainty assessed using a Fieller interval derived from the shared-design covariance of the two endpoint slopes.

In a separate exploratory cell-level analysis, single-cell observations were centered and scaled within well, and the within-well association between cell-level ATeam and morphology metric was contrasted between each compound/regime and the main DMSO vehicle reference using well-clustered standard errors; this cell-level comparison did not alter the well-level L1-L3 classification. The hierarchy is descriptive and does not imply causal direction between mitochondrial function and morphology. Statistical analyses were performed in R using base R stats and data.table; related R figure-generation scripts used ggplot2, ggnewscale, ggrepel, and svglite. The dotplot was rendered using a Python/matplotlib script. The initial analysis framework, including R-script development, exploratory statistical design, and provenance documentation, was started using Claude.ai with Claude Opus 4.8. Subsequent code modification, analysis reruns, figure/table generation, result interpretation, and manuscript-methods copyediting were completed using OpenAI Codex with GPT-5.5 and were human verified. Analysis codes and skills are available at https://github.com/gerencserlab/Co-Response.

### Statistics

Pairwise and basic multiple comparisons were performed as given in text using GraphPad Prism 11. Bayesian linear regression was used to analyze treatment-associated assay responses, with model structure matched to the experimental design: respirometry data were analyzed in Python using Pandas, NumPy, and SciPy as Gaussian percent-change outcomes with treatment-by-substrate effects and subject blocking terms for repeated donor measurements, whereas potentiometric membrane-potential data were analyzed in Wolfram Mathematica 13-14 using ResourceFunction["BayesianLinearRegression"] after standardization of the response and covariates, with encoded dispense and microscope conditions plus age, HbA1c, BMI, and sex as predictors and no blocking. For both analyses, posterior coefficient distributions were summarized by point estimates and 95% credible intervals (CrI), with effects interpreted as credibly different from zero when the CrI excluded zero; respirometry treatment-substrate tests were additionally controlled for multiple comparisons using false-discovery-rate (FDR) adjustment using the Benjamini-Hochberg (BH) procedure.

For claims on equivalent coefficients of variation (CoV), equivalence was assessed using two one-sided Welch tests (TOST) with a symmetric equivalence margin of ±3 or ±5 CoV percentage points and α = 0.05. Equivalence was assessed after neither Welch t-test nor one-sample Wilcoxon signed-rank test (where appropriate) detected a difference.

Dispersion of assay window metrics and variances was determined by bootstrapping in Python, treating microplates as replicates. Data were adjusted for multiple testing as indicated in text, showing either BH FDR values or Holm correction for multiple testing (in cases of 2-4 groups).

### Software and Code Availability

PickliPy requires Python ≥3.9 and standard scientific computing libraries including Pandas, NumPy, and openpyxl. All scripts are platform-independent and tested across Windows and macOS environments. PickliPy.QC is a companion tool for post-execution validation that parses Echo print and survey XML log files, matches transfers to compound library data, and generates consolidated quality control reports. PickliPy.QC is available both as Python source code and as a standalone Windows executable requiring no programming experience or software dependencies. Source code, documentation, templates, and executables are available at https://github.com/gerencserlab/PickliPy.git.

Input workbooks were created and edited in Microsoft Excel. Metadata recording assay-well associations are saved by PickliPy and are compatible with Image Analyst MKII workflows analyzing image data from Revvity Harmony or with DataWarrior. Generated plate and concentration maps may be copied as assay layouts into Harmony. Picklist and accompanying inventory and process files were used with Revvity Plate::Works^TM^ (version 6.30) scheduling software and Beckman Coulter Echo software (version 3.2.3). Image Analyst MKII pipelines and corresponding Python wrappers are available at the above provided repositories.

### AI-assisted computational analysis

Parts of the computational analysis workflow were developed and documented with agentic large-language-model assistance under human supervision. This includes Claude.ai with Claude Opus 4.8 and OpenAI Codex with GPT-5.5. The AI agents were used to generate and revise analysis code, inspect outputs, suggest statistical summaries, and draft documentation; they were not listed as authors and did not make independent scientific conclusions. AI agents were also used for literature search, analyzing instrument documentation, manuscript copyediting, and translating Wolfram Language code to Python and writing software code for image analysis. All code, commands, outputs, statistical decisions, figure files, and manuscript text were reviewed, edited, and approved by the human authors, who take full responsibility for the integrity, reproducibility, and interpretation of the work.

## Supplementary Material

**Table S1.**
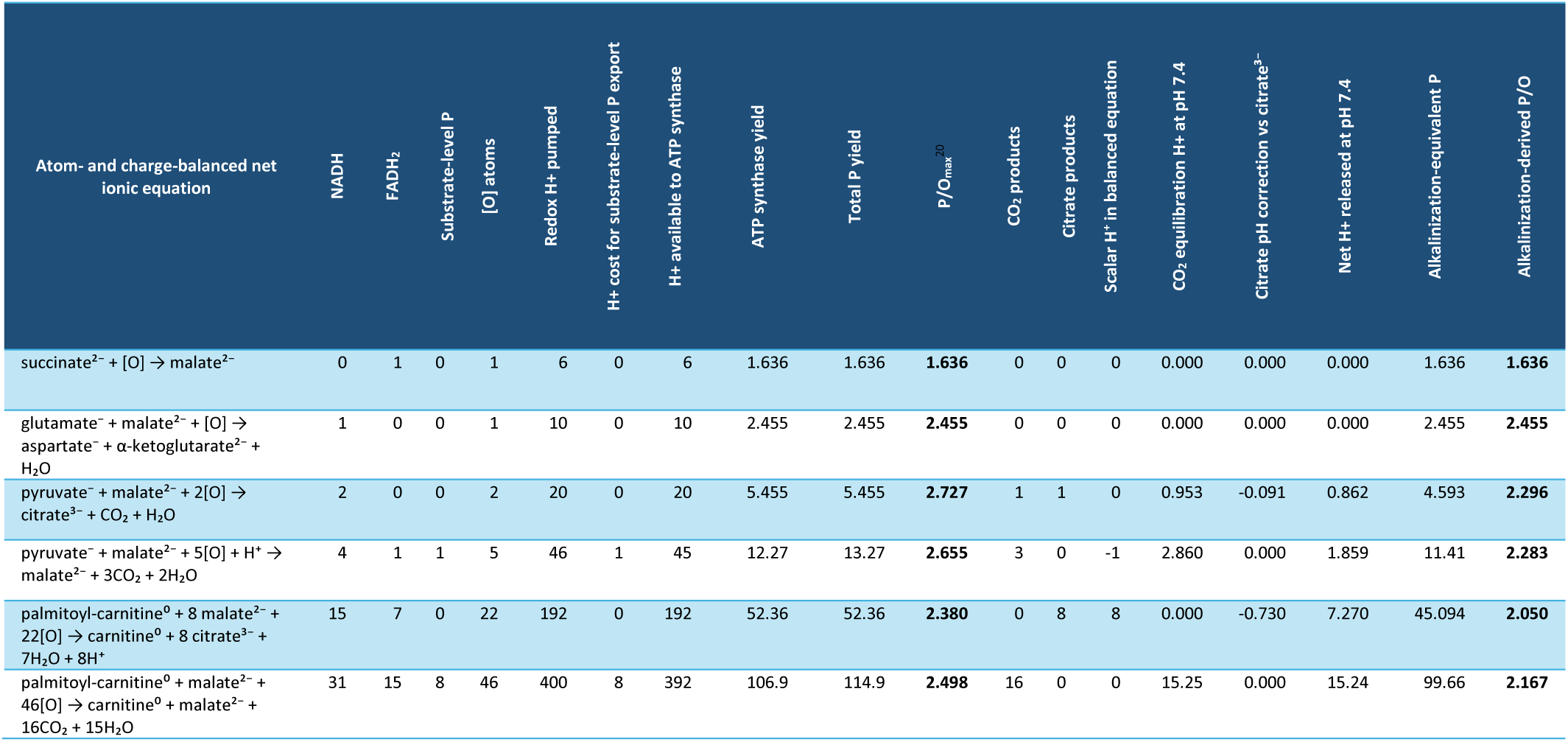
Calculation of expected alkalinization-derived P/O values. . The calculations extend the assumptions applied by Mookerjee et al.^20^ by considering net acid and base formation and the substrate combination of palmitoyl-carnitine + malate with complete or incomplete oxidation.

**Table S2.** Consumables, listing microplates used in the study.

| Manufacturer | Description | Catalog # |
| --- | --- | --- |
| Beckman Coulter, Palatine, IL | Echo® Qualified 384-Well Polypropylene 2.0 Microplate, Clear, Flat Bottom | 01-14615 |
| Beckman Coulter, Palatine, IL | Echo® Qualified 384-well Cyclic Olefin Copolymer Source Microplate, Low Dead Volume, Clear | 001-16128 |
| Cellvis, Mountain View, CA | 384-well glass-bottom plate with high performance #1.5 cover glass | P384-1.5H-N |
| VWR International | MICROPLATE 96-WELL CLEAR CS100 (Corning #3363) | 29444-102 |
| Merck KGaA - Sigma-Aldrich, St. Louis, MO | CORNING(R) HIGH CONTENT IMAGING PLATE #4580 | CLS4580 |
| Agilent Technologies, Santa Clara, CA | Seahorse XFe96/XF Pro FluxPak | 103792-100 |

**Table S3.** Post-mortem pancreatic islet donor data. Table shows islet centers and identifiers associated with each islet preparation, islet donor clinical values and assay-related information. FCCP or Atpenin A5 dose-response experiments were scheduled based on prioritizing other (not shown) experiments. For benchmarking ADE with FCCP dose response the last 24 donors were included with 50% of them analyzed using ADE. For Atpenin A5 dose-response experiments all historical data were included in the analysis. Values in the ΔψM (mV) columns are either mean ± SE of 2–3 experimental replicates or mean of 2 technical replicates. The depolarizing effect of FCCP was measured in the presence of 16 mM glucose. Blank cells indicate measurements that were not performed.

| ID | Islet Center | Age (y) | Sex | BMI | HbA<sub>1c</sub> | T2D Status | Dispense method | Microscope | Included to FCCP dose-response | Included to Atpenin A5 dose-response | $\Delta\psi_M$ depolarization @ 100 nM FCCP (mV) | $\Delta\psi_M$ depolarization @ 300 nM FCCP (mV) | EC50 for Atpenin A5 (μM) |
| --- | --- | --- | --- | --- | --- | --- | --- | --- | --- | --- | --- | --- | --- |
| HP-17300 | Prodo | 35 | M | 27.7 | 6.3 | T2D | Manual | 1 | FALSE | TRUE | 35.1±1.7 | 63.8±2.4 | 0.0904 |
| HP-17342 | Prodo | 38 | M | 30 | 5.8 | normal | Manual | 1 | FALSE | TRUE | 47.6±3.1 | 64±4 | 0.307 |
| HP-18275 | Prodo | 30 | F | 40.1 | 6.5 | T2D | Manual | 1 | FALSE | TRUE | 48±4 | 79±4 | 0.0793 |
| HP-18320 | Prodo | 61 | M | 27.4 | 7.1 | T2D | Manual | 1 | FALSE | TRUE | 41±4 | 79±8 | 0.497 |
| HP-18330 | Prodo | 53 | F | 26.3 | 5.4 | normal | Manual | 1 | FALSE | TRUE | 51.3 |  | 0.275 |
| HP-19017 | Prodo | 45 | F | 36.1 | 5.7 | normal | Manual | 1 | FALSE | TRUE | 43±4 |  | 0.226 |
| HP-19051 | Prodo | 53 | M | 30.1 | 7.8 | T2D | Manual | 1 | FALSE | TRUE | 20±4 | 46±6 | 0.201 |
| HP-19242 | Prodo | 55 | M | 28.4 | 5.6 | normal | Manual | 1 | TRUE | FALSE | 37.8±1.7 | 64.6±1.7 |  |
| rHIP-136 | UCSF | 53 | M | 21.3 | 5.4 | normal | Manual | 1 | TRUE | FALSE | 33±8 | 65±10 |  |
| HP-19276 | Prodo | 37 | M | 24.2 | 5.8 | normal | Manual | 1 | TRUE | FALSE | 47.4±1.4 | 63±4 |  |
| HP-19283 | Prodo | 43 | F | 31.9 | 5.7 | normal | Manual | 1 | TRUE | FALSE | 41.6±2.8 | 64±4 |  |
| rHIP-137 | UCSF | 35 | F | 33.3 | 5 | normal | Manual | 1 | TRUE | FALSE | 29.7 | 58 |  |
| HP-19324 | Prodo | 43 | F | 23.2 | 5.4 | normal | Manual | 2 | FALSE | TRUE | 44±4 |  | 0.229 |
| HP-19342 | Prodo | 64 | M | 21 | 5.5 | normal | Manual | 2 | TRUE | TRUE | 36.3±2.7 | 61±4 | 0.179 |
| HP-20019 | Prodo | 53 | M | 25.8 | 6.7 | T2D | Manual | 2 | TRUE | FALSE | 43±4 | 71±4 |  |
| HP-20021 | Prodo | 52 | M | 29.6 | 5.4 | normal | Manual | 2 | TRUE | TRUE | 52.7 | 88.1 | 0.178 |
| HP-21247 | Prodo | 53 | M | 32.2 | 5.9 | normal | Manual | 2 | TRUE | TRUE | 44±4 | 89±10 | 0.366 |
| HP-21263 | Prodo | 62 | M | 34.6 | 6.8 | T2D | Manual | 2 | TRUE | TRUE | 34.4±3.0 | 68.6±1.3 | 0.29 |
| rHIP-149 | UCSF | 40 | F | 23.7 | 5.8 | normal | Manual | 2 | TRUE | TRUE | 29±6 | 60.4±3.5 | 0.193 |
| HP-22159 | Prodo | 47 | M | 29.7 | 5.2 | normal | Manual | 2 | TRUE | TRUE | 27.7 | 56 | 0.0977 |
| HP-24220 | Prodo | 47 | M | 29.6 | 5.2 | normal | ADE | 1 + 2 | TRUE | FALSE | 35.5±2.2 | 59±4 |  |
| HP-24222 | Prodo | 43 | M | 27 | 5.5 | normal | ADE | 2 | TRUE | FALSE | 30±4 | 53.4±1.5 |  |
| HP-24254 | Prodo | 57 | M | 28.8 | 5.9 | normal | ADE | 2 | TRUE | FALSE | 40.9±1.7 | 85±6 |  |
| HP-24303 | Prodo | 49 | M | 24.5 | 5 | normal | ADE | 1 | TRUE | FALSE | 33.1 | 74.4 |  |
| HP-25063 | Prodo | 59 | M | 24 | 5.7 | normal | ADE | 1 | TRUE | TRUE | 26.4 | 65 | 0.087 |
| HP-25086 | Prodo | 47 | M | 26.2 | 5.4 | normal | ADE | 1 | TRUE | TRUE | 37±4 | 75±7 | 0.137 |
| HP-25113 | Prodo | 56 | M | 28.3 | 5.3 | normal | ADE | 1 | TRUE | FALSE | 27±6 | 62±7 |  |
| HP-25135 | Prodo | 40 | F | 25 | 5 | normal | ADE | 1 | TRUE | TRUE | 47±17 | 78±23 | 0.0579 |
| HP-25227 | Prodo | 54 | M | 31.3 | 5.3 | normal | ADE | 1 | TRUE | TRUE | 39.5±2.2 | 73±4 | 0.117 |
| HP-25276 | Prodo | 41 | F | 19.8 | 5.1 | normal | ADE | 1 | TRUE | TRUE | 30.5±1.7 | 78.8±1.7 | 0.0585 |
| SAMN54469755 | IIDP (Prodo) | 61 | M | 29.3 | 5.4 | normal | ADE | 1 | TRUE | FALSE | 29±9 | 67±11 |  |
| SAMN54754376 | IIDP (Imagine Pharma) | 57 | M | 30.4 | 8.4 | T2D | ADE | 1 | TRUE | TRUE | 20±5 | 54±7 | 0.475 |

**Fig. S1.**
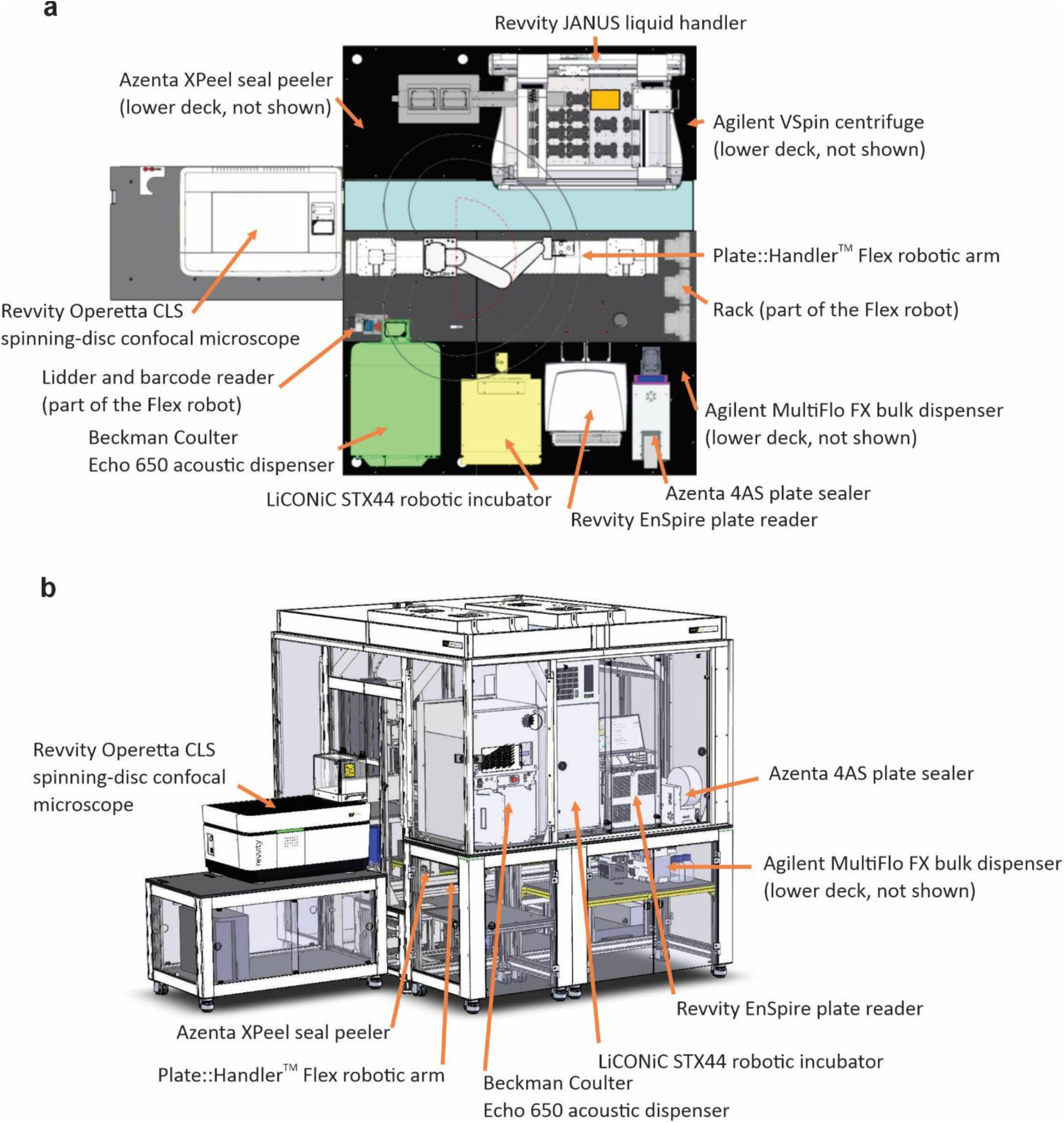
**Robotic workcell configuration** shown as **a)** top view and as **b)** 3D rendering at design stage.

**Fig. S2.**
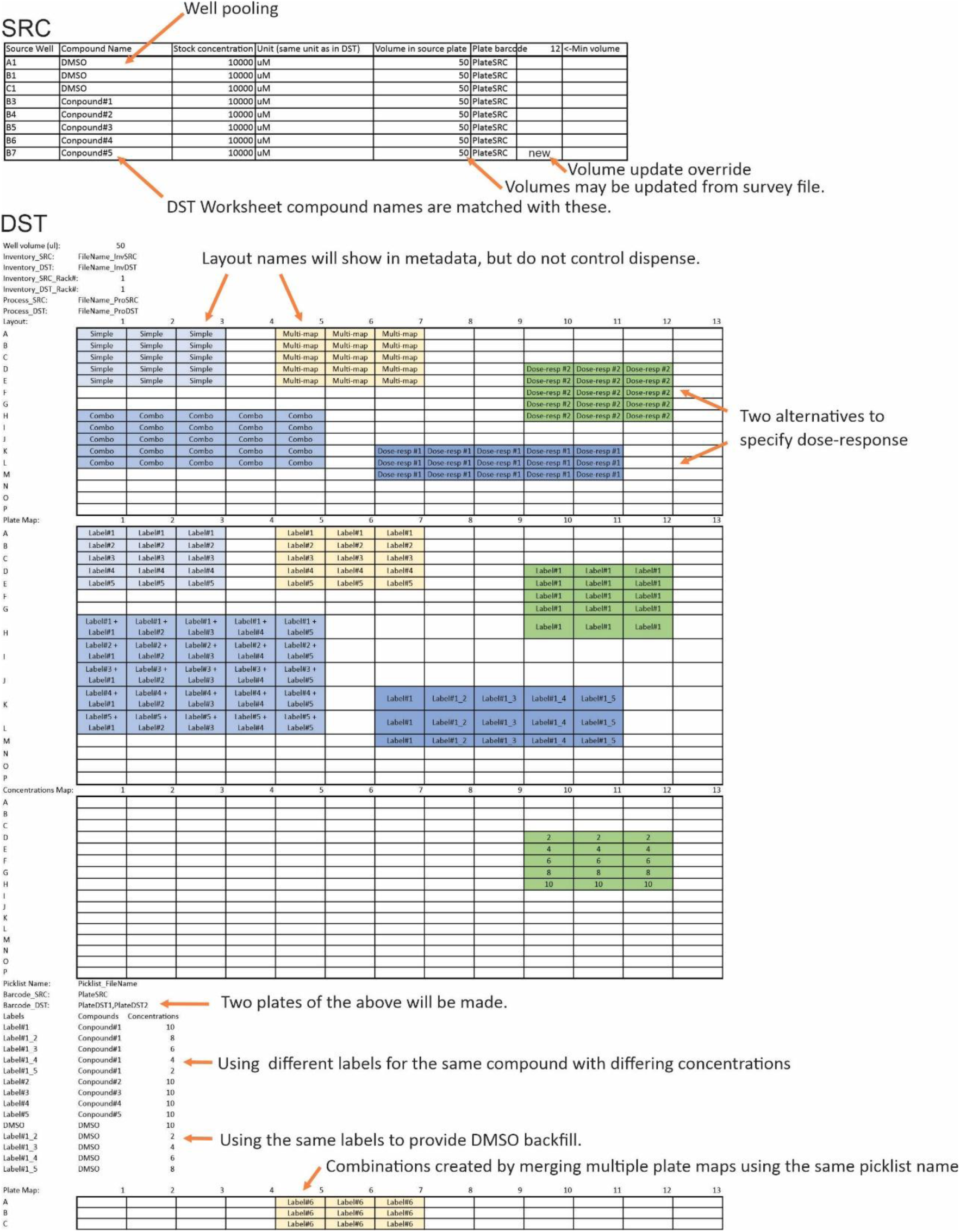
Example design file for PickliPy.Assay. Different DST worksheet layout labels indicate different design strategies that can be all combined on a single plate.

**Fig. S3.**
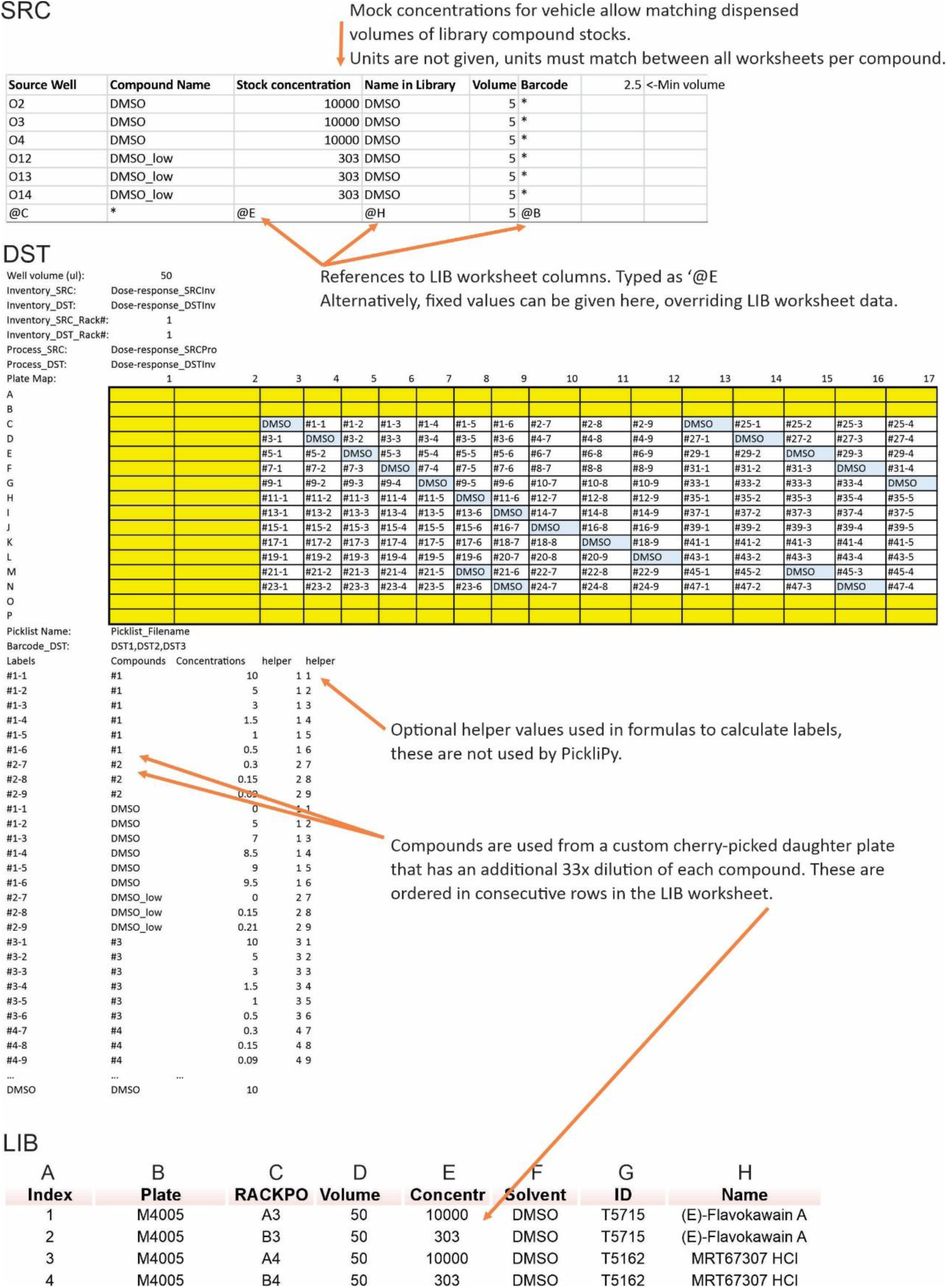
Example design file for PickliPy.Screen. Layout corresponds to Fig. 1e.

**Fig. S4.**
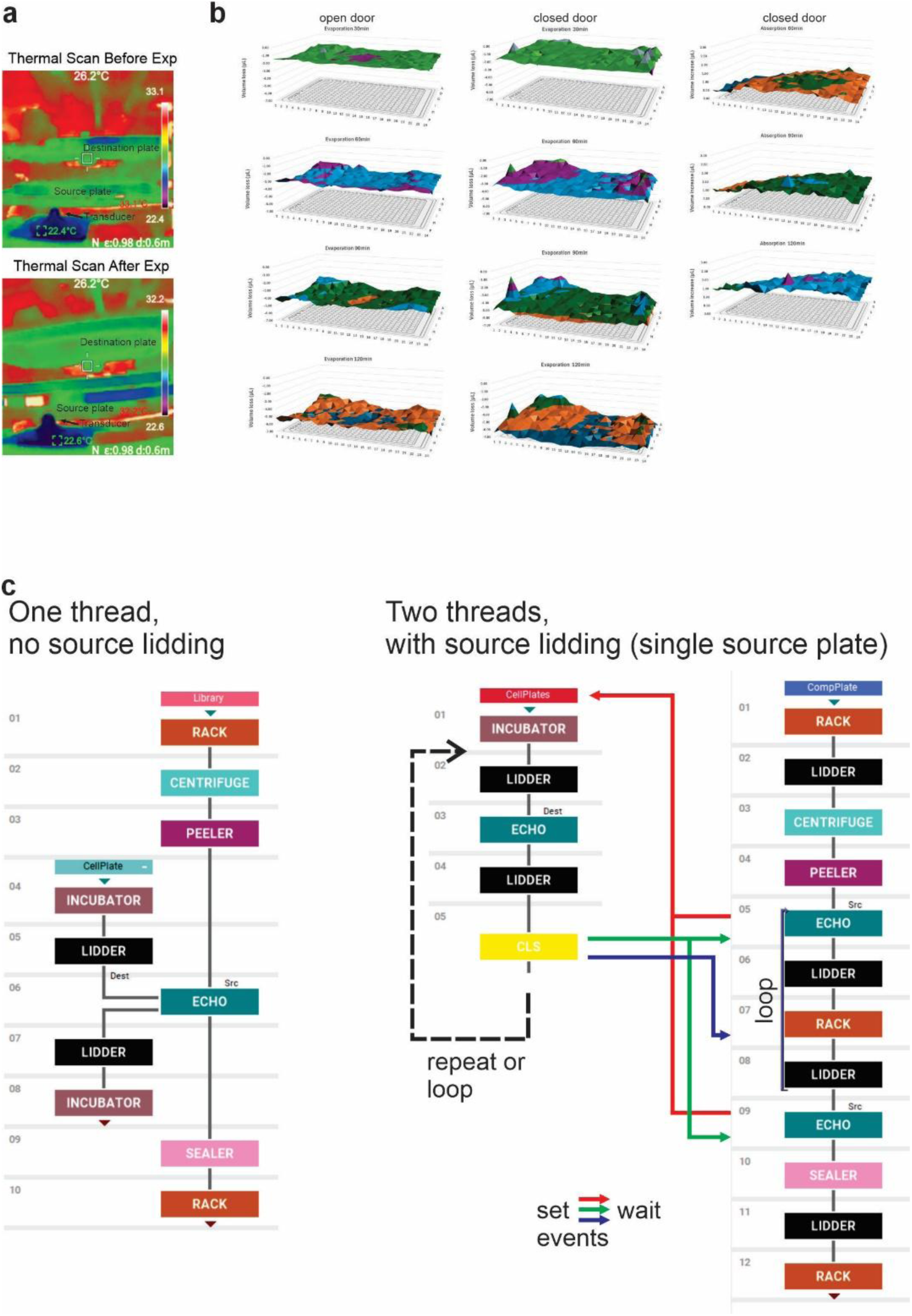
Evaporation and moisture absorption during ADE. **a)** Thermal imaging of the dispense chamber of the Echo 650. **b)** Evaporation and moisture absorption data are presented as a 3D surface plot to show the change in liquid levels across the plate over time, with dispense chamber door opened or closed during the wait period. The door is normally closed during dispense but opened in wait periods, e.g. during time courses. **c)** Revvity Plate::Works^TM^ assay diagrams showing a simple ADE dispense paradigm, compatible with library screening (left) and a two-thread design allowing lidding of the source plate in the wait periods, compatible with single source plate time courses (right).

**Fig. S5.**
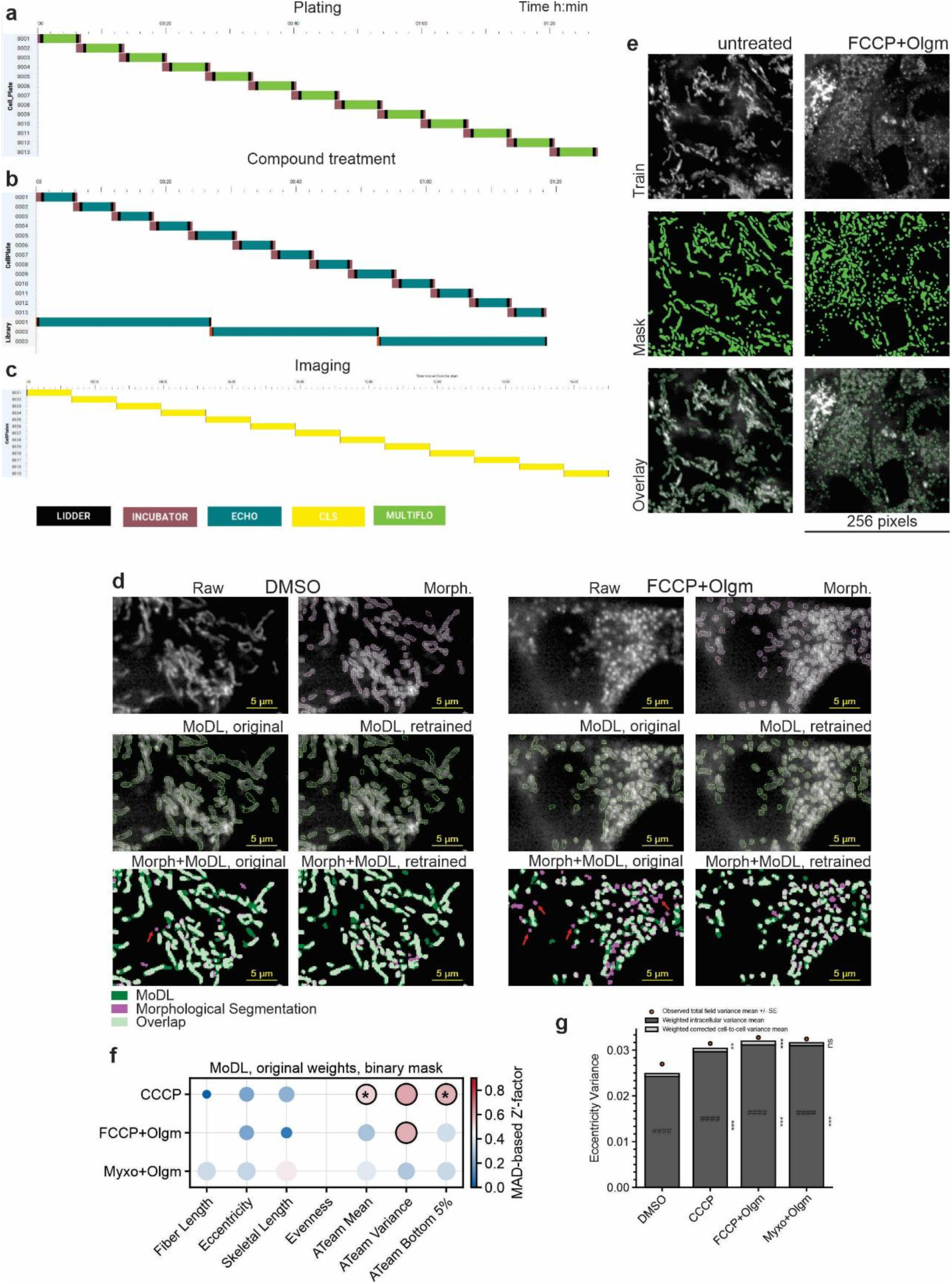
Small-molecule library screening with reformatting and three doses per plate for modulators of mitochondrial [ATP] and morphology using PickliPy.Screen. **a)** Gantt chart of cell plating using the MultiFlo FX bulk dispenser. **b)** Gantt chart of compound treatment using the Echo 650. **c)** Gantt chart of imaging using the Operetta CLS. **d)** Comparison of morphological segmentation with MoDL segmentation using pretrained and in-dataset retrained network weights. The inputs were mito-ATeam fluorescence images (Raw), in control (DMSO) and in FCCP+Olgm (4 µM + 2 µg/mL) conditions. Overlays show pretrained and retrained MoDL segmentation in green and morphological segmentation in purple. Red arrows point to fragmented mitochondria captured by morphological segmentation (purple) but not by the pretrained network. Data correspond to Fig. 5k,l. **e)** Representative training and curated reference masks (serving as ground truth images) for MoDL retraining, showing a quarter of the training image. The mask was generated as shown in d) “Morph+MoDL, original” by adding standalone objects shown in purple to the green MoDL mask. **f)** MAD-based Z′-factor (*Z*′_MAD_) of screening with metrics illustrated in Fig. 5m and corresponding to Fig. 5n, comparing the positive controls to DMSO. Here, *Z*′_MAD_ was obtained after image analysis using the original network weights of MoDL, with 4× upscaling to better match training and data set resolutions, and directly using the binary mask produced by the model without post-processing. Dots indicate *Z*′_MAD_ > 0; black outlines, *Z*′_MAD_ > 0.5, *, 95% CI of *Z*′_MAD_ above 0.3. **g)** Variance hierarchy decomposition to distinguish contribution of cell-to-cell and intracellular variation (stacked bars) to the total, view field level variance (data points) of eccentricity (corresponding to Fig. 5o, p). **, ***, BH FDR q ≤ 0.01 and 0.001, respectively, compared to the same metric in DMSO. ### and #### indicate the larger component at BH FDR q ≤ 0.001 and 0.0001, respectively, within each stack.

**Fig. S6.**
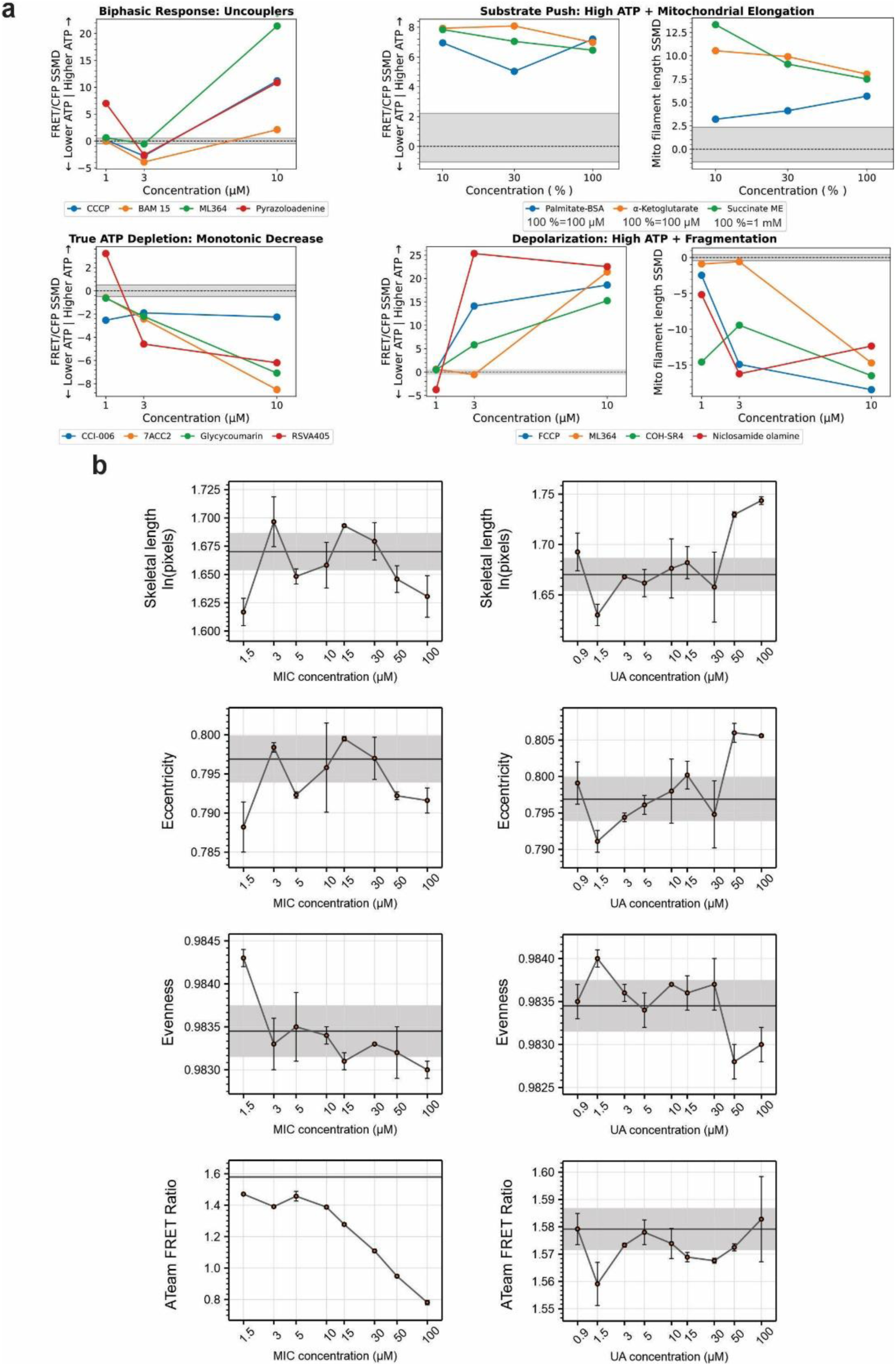
Dose-response experiments in the mito-ATeam HEK293 cell screen and with cherry-picked library compounds. **a)** Distinct dose-response patterns observed in the main screen. Labels indicate known properties of exemplified compounds or observations in the fiber length data (not shown). Each data point represents one primary-screen measurement. **b)** Dose-response relationships showing mean ± SE of n = 3 plate replicates with randomized layouts in the screen follow-up, showing effects of MIC and UA.

**Fig. S7.**
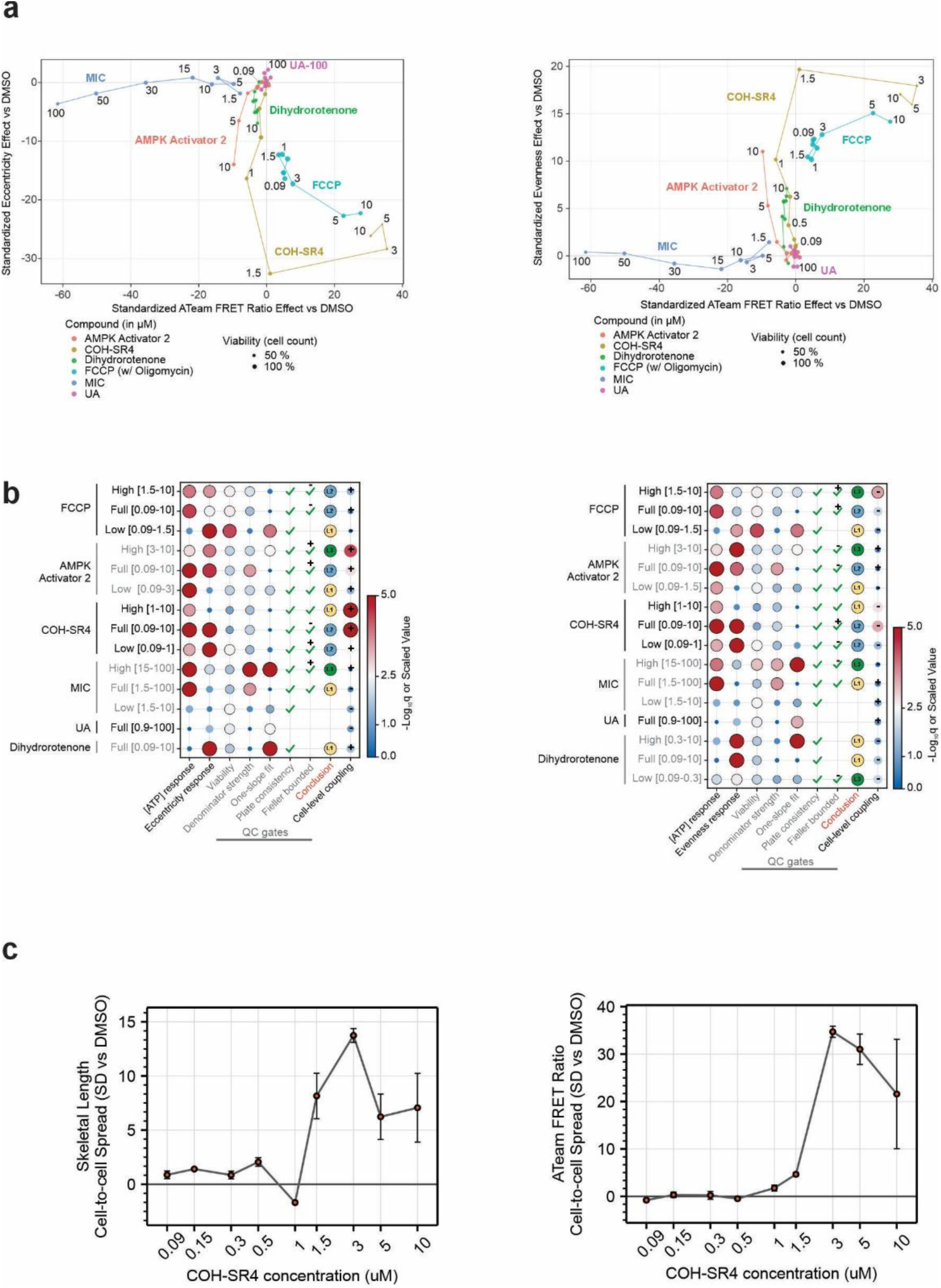
Dose-response co-response hierarchy for ATeam FRET ratio and morphological metrics in HEK293 cells. **a)** Function-morphology effect-space trajectories of DMSO-standardized ATeam FRET ratio as a function of mitochondrial eccentricity or mitochondrial thickness evenness (calculated as normalized Shannon effective-number of distance values from the skeleton pixels to the boundary). Lines connect increasing concentrations, providing a dose-dependent path through the function-morphology space. Distinct trajectories indicate that eccentricity or thickness evenness did not simply mirror the ATeam FRET ratio across perturbations; instead, the relationship between functional and morphological phenotypes varied by compound and dose range. **b)** Dose-response co-response hierarchy for ATeam FRET ratio and eccentricity or thickness evenness. Rows summarize full-range, auto-selected breakpoint-defined high and low concentration windows for each compound. Columns show the evidence used to classify morpho-functional coupling. Black outlines mark accepted gates or conclusions (FDR q < 0.05 for responses, for others see Methods). The Fieller bounded column indicates that an interval for effect-dose slope ratios can be robustly defined, with + or – indicating that there is a significant coordinated or anti-coordinated dose-response. The Conclusion column reports the hierarchy level: L1, single-endpoint or dissociated response; L2, dose-associated co-response; and L3, proportional local coupling. The cell-level coupling column indicates significance level and direction of the compound-induced change. Responses and coupling are provided in -log_10_q values capped at 5. c) Standardized spread of skeletal length and ATeam FRET ratio as a function of COH-SR4 concentration, indicating compound-induced cell-to-cell heterogeneity. Data points are mean ± SE of n = 3 plate replicates.

**Fig. S8.**
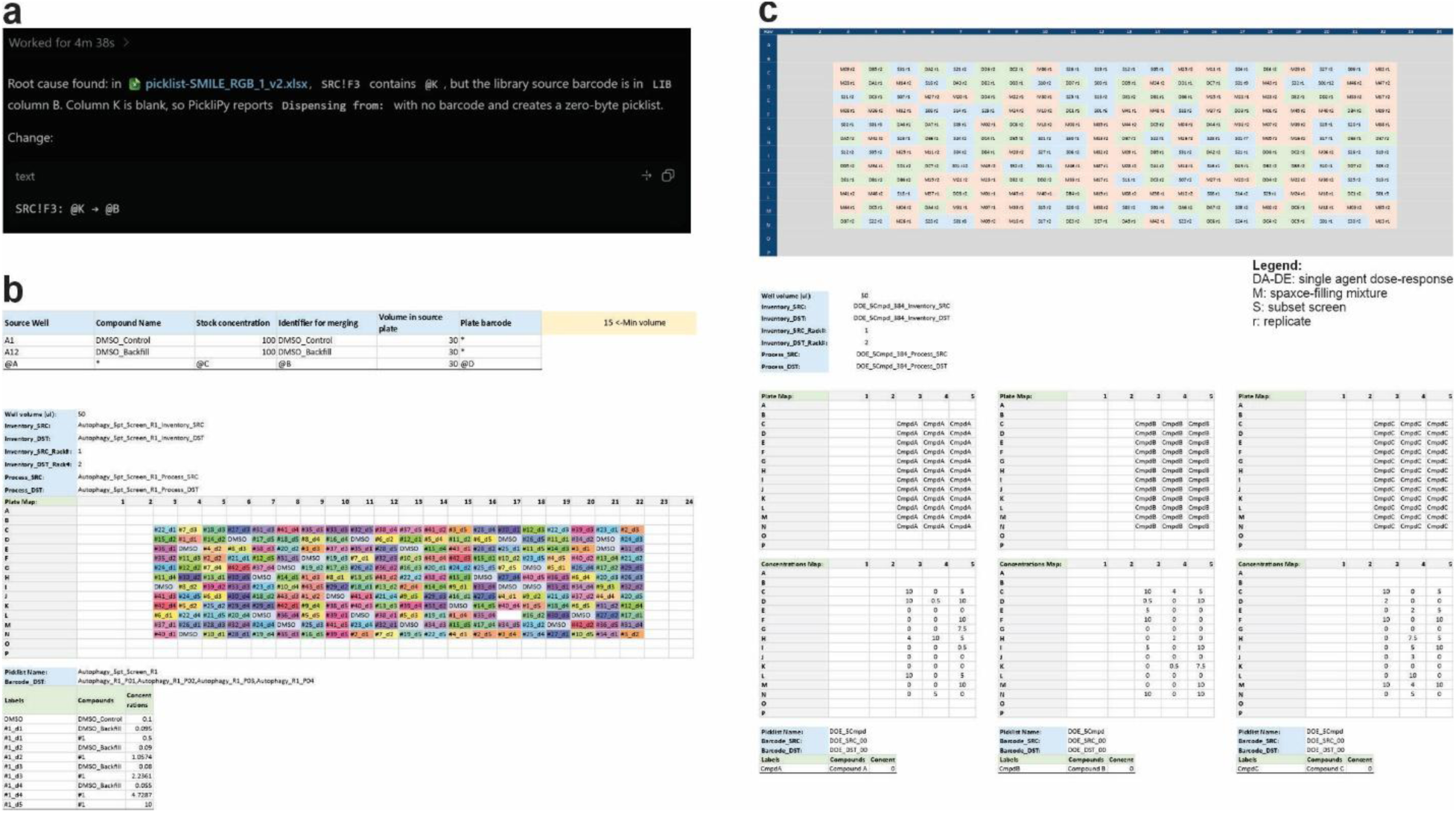
PickliPy used as an agentic tool. **a)** Agentic troubleshooting of user error. Prompt: Use picklipy-excel-design-skill to troubleshoot a design file using PickliPy.Screen. The problem file is test_smile\picklist-SMILE_RGB_1_v2.xlsx. I want to dispense a pattern using one of 5 solutions into each well of a 384-well plate, but I don’t get any dispense events. **b)** Agentic generation of a dose-response experiment using a library subset. Prompt: Use picklipy-excel-design-skill to make a design file and run PickliPy.Screen for a 5-point dose response using only compounds from the library that mention autophagy. Add 24 DMSO controls to each plate, in evenly-distributed random positions. Add DMSO backfill, so the vehicle concentration is always 1:1000. Use center 240 wells only. The library file is in test_screen\L5300-Mitochondria-Targeted Compound Library-950cpds.sdf. Work in that folder. Use concentrations between 0.5 and 10 uM in log steps. The stocks are 10 mM. Make 3 randomized replicates of each plate. Add cell coloring to the plate map, unique for each compound on that plate. **c)** Agentic generation of a highly complex combinatorial plate design. Top: conditions layout, Bottom: DST worksheet showing only first 5 columns, and first 3 compounds. Prompt: *Using picklipy-excel-design-skill create a design file using PickliPy.Assay for a design-of-experiment (DOE) style assay on a single destination plate. In the source plate I have 5 compounds in wells A1-E1, 10 mM stocks. Call them Compound A-E. Use minimum 0.5 uM and maximum 10 uM concentrations, in 50 ul assay volume. The experiment must be able to determine that in what combination are these compounds the most effective on a univariate, continuous readout. Use 384-well format, with center 240 wells in use. Explain the strategy and advise on downstream data analysis*.

